# Exploring the roles of trophic mode and microbial prey size in grazing pathways of tropical oligotrophic waters of the eastern Indian Ocean

**DOI:** 10.1101/2025.09.02.673656

**Authors:** Claudia Traboni, Grace F. Cawley, Karen E. Selph, Michael R. Landry, Moira Décima

## Abstract

Prey removal incubations were conducted in the Argo Basin (eastern Indian Ocean) to investigate the trophic ecology of the zooplankton community supporting Southern Bluefin Tuna larvae. Grazing and selectivity were evaluated considering prey trophic mode and size as food quality descriptors in copepod diets and compared with microzooplankton grazing. Copepod ingestion of eukaryotes ranged from 3.4 to 138 ng carbon (C) ind^-1^ d^-1^. Diet was dominated by mixotrophic (5-89%) and heterotrophic (0-84%) prey, with autotrophs contributing 2-17%. Nanoplankton provided the highest C intake to copepods (62-99%) rather than more abundant picoplankton (0.8-38%). No measurable reduction in chlorophyll a (Chl*a)* concentration occurred from copepod grazing through food removal, suggesting a possible trophic cascade, although gut content revealed ingestion of 0.8 µg Chl*a* ind^-1^ d^-1^. Copepods showed moderate selection for picoplankton (E=0.3) over nanoplankton, perhaps due to faster consumption of nanoplankton by microzooplankton or ingestion of picoplankton aggregates. Ingestion of ^15^N (nitrogen)-labelled prey indicated copepod consumption of protistan consumers or small metazoans. We found significantly greater (∼5-fold) copepod N consumption when using 1-2 µm ^15^N-*Synechococcus*, compared to <1 µm sized ^15^N-*Prochlorococcus*. Microzooplankton grazing on eukaryotes (0.07–2.5 d^-1^) and prokaryotes (0.3–2.1 d^-1^) greatly exceeded copepod grazing. Microzooplankton diet consisted mainly of heterotrophs (25-59%) and mixotrophs (13-41%), with lower autotrophic contribution (12-33%) of more nano- (95-98%) than picophytoplankton (2-5%) ingested. Overall, microzooplankton removed most of daily production (111%) in contrast to 7% for copepods. Our findings indicate that mixotrophy, intraguild grazing and nutrient channeling support the food web in this oligotrophic region.

## 1. Introduction

Copepods feed on a wide range of protistan prey varying in taxonomy, size, mobility, trophic mode and biochemical composition (Saiz and Calbet, 2011). The size ratio between prey and predator is a master trait determining capture and ingestion probability (Hansen et al., 1994), but trophic mode can also be an important factor affecting nutritional quality of copepod prey (Traboni et al., 2021). Autotrophic phytoplankton depend on inorganic nutrients and sunlight to sustain their photosynthetic metabolism and accumulate carbon-rich biomass, whereas heterotrophic protozooplankton are obligate consumers of organic matter that are more enriched in nitrogen and phosphorus relative to carbon. Mixotrophic protists (mixoplankton) combine both trophic modes (Flynn et al., 2019; Stoecker et al., 2017) and are characterized by intermediate nutritional composition. Especially under conditions where dissolved concentrations of nutrients are very low, the ability of mixotrophs to acquire nutrients from phagotrophy can improve food quality by reducing prey–predator stoichiometric mismatch (Schenone et al., 2024) and enhancing nutrient transfer efficiency (Ward and Follows, 2016). Laboratory experiments have demonstrated that copepods feed abundantly on mixotrophs (Jeong et al., 2010; Ptacnik et al., 2004) and are not negatively affected by consumption of mixotrophs compared to other trophic modes (Traboni et al., 2020). However, only a few field studies consider this food-web interaction (Castellani et al., 2005; Fileman et al., 2007; Jonsson and Tiselius, 1990), likely due to methodological challenges in distinguishing mixotrophs in natural assemblages.

Mixotrophy is a prevalent adaptive strategy among planktonic protists of oligotrophic regions, overcoming resource limitation of photo- and phagotrophy specialists (Anschütz et al., 2024; Edwards, 2019) by using photosynthesis for growth when nutrients allow and phagotrophy when nutrients/light are scarce (Stoecker et al., 2017). The mixotrophic strategy has been reported worldwide in oligotrophic regions of the North Atlantic (Hartmann et al., 2012; Millette et al., 2016), East China Sea (Li et al., 2024), Sargasso Sea (Arenovski et al., 1995) and Mediterranean Sea (Unrein et al., 2014). Of particular relevance to the current study, Raes et al. (2022) found an 18S-rRNA diversity peak for mixotrophs in high-temperature low-nutrient tropical waters of the eastern Indian Ocean (IO). Elevated food-web transfer efficiency associated with mixotrophy was also invoked to explain especially high zooplankton grazing on chlorophyll-containing protists in this region (Landry et al., 2020a). In oligotrophic systems, phytoplankton communities are typically dominated by small autotrophs, while larger phytoplankton are scarce and episodic. Microzooplankton communities (<200 µm) are dominated by dinoflagellates, ciliates and some metazoan larvae closely coupled to the picoplankton prey base (Sherr and Sherr, 2002). Mesozooplankton assemblages (>200 µm) are usually dominated by copepods, which rely on nano-microzooplankton and occasional pulses of relatively large phytoplankton (Calbet, 2008; Saiz and Calbet, 2011). As a result, grazing pressure in oligotrophic waters follows a size-structured pattern, with microzooplankton dominating consumption of prey <20 µm, and copepods targeting larger prey or exerting indirect control via predation on nano-microzooplankton, at times overlapping with microzooplankton in nanoplankton ingestion.

In January-March 2022, the BLOOFINZ-IO (Bluefin Larvae in Oligotrophic Ocean Foodwebs, Investigations of Nutrients to Zooplankton – Indian Ocean) expedition investigated the physical, biogeochemical and food web characteristics of waters overlying the Argo Abyssal Plain (hereafter, Argo Basin), a 5000-m deep oceanic basin directly downstream of the Indonesian Throughflow (ITF), where the tropical surface water mass from the western Pacific first enters the eastern IO (Woo and Pattiaratchi, 2008). This is also the only known global spawning region for Southern Bluefin Tuna, *Thunnus maccoyii* (Davis et al., 1991), which underlies the BLOOFINZ goal of understanding details of the food web supporting feeding, growth and survival of the species through its vulnerable larval stages. As part of this effort, we conducted experimental incubations to explore the roles of size structure and trophic mode in grazing pathways to microzooplankton and copepods, with a focus on mixo-heterotrophy. We hypothesized that mixotrophs and heterotrophs would dominate the community composition in this oligotrophic region and ii) that mixotrophs would be a high fraction of the copepod diet accordingly. To test these hypotheses, flow cytometry and acidic vacuole staining were applied to characterize prey communities (abundance of cells, size and trophic mode), food removal incubations served to quantify grazing rates and selectivity, and isotope labelling was also included to trace the main nutrient pathway from lower food web to copepods and as an indication of copepod dietary origin.

## 2. Materials and methods

### 2.1. Grazing experiments: study area and timeline of the incubations

Zooplankton grazing experiments were performed aboard R/V *Roger Revelle* during BLOOFINZ-IO cruise RR2201, with sampling taking place between February 4^th^ and March 2^nd^ 2022, off NW Australia (Argo Basin) (Fig. 1A, B). During the cruise, daily CTD casts (Sea Bird Scientific 911) were conducted to monitor vertical profiles of abiotic (temperature, salinity, irradiance, oxygen) and biotic features (i.e., chlorophyll a (Chl*a*), microbial abundance and biomass) of the water column (Table 1), as well as to collect water for experiments through 10-L Niskin bottles mounted on a 12-place rosette. Grazing incubations were mostly conducted during multi-day experiments (hereafter “cycles”) following a satellite-tracked drifter (Landry et al., 2009; Landry et al., 2025). Nine grazing experiments were conducted in total (Exps 1-9). The first seven experiments were performed during Cycles 1-4, and two final grazing incubations were conducted during transits along the return leg of the cruise (Table 1; Fig. 1B). Each experiment consisted of two incubation phases: a 24-h incubation with the microzooplankton community alone (“Microzoo” incubation, T_0_-T_24_) followed by a 12-h incubation with added copepods (“Microzoo+copepod” incubation, T_24_-T_36_) (Fig. 2, Table 2).

**Table 1.** General spatio-temporal details of the grazing experiments and main parameters relative to seawater sampled through Niskin bottles during evening CTD casts and zooplankton live net sampling. Abbreviations: Exp: experiment, Lat: latitude, Long: longitude, Temp: temperature, Oxy: dissolved oxygen, FChl*a*: chlorophyll *a* from fluorescence; Zoo: zooplankton.

|  |  |  |  | CTD cast – Niskin bottle seawater sampling |  |  |  |  |  |  | Zoo sampling |
| --- | --- | --- | --- | --- | --- | --- | --- | --- | --- | --- | --- |
|  |  |  |  | Lat | Long | Depth | Salinity | Temp | Oxy | FChla | Depth |
| Date | Cycle / Station | Cycle day | Exp | °N | °E | (m) | | (°C) | ($\mu\text{mol L}^{-1}$) | ($\mu\text{g L}^{-1}$) | (m) |
| 04 Feb. | 1 | 1 | 1 | -15.37 | 114.43 | 30 | 34.53 | 28.61 | 190.7 | 0.118 | 48 |
| 09 Feb. | 2 | 1 | 2 | -16.74 | 115.69 | 30 | 34.93 | 27.93 | 193.5 | 0.118 | 38 |
| 12 Feb. | 2 | 3 | 3 | -16.99 | 116.05 | 30 | 34.97 | 28.09 | 192.6 | 0.108 | 38 |
| 15 Feb. | 3 | 1 | 4 | -15.98 | 115.83 | 28 | 34.66 | 28.63 | 194.8 | 0.125 | 41 |
| 17 Feb. | 3 | 3 | 5 | -15.81 | 115.82 | 30 | 34.56 | 28.01 | 195.3 | 0.143 | 38 |
| 20 Feb. | 4 | 1 | 6 | -15.89 | 118.14 | 30 | 34.52 | 29.18 | 193.5 | 0.144 | 60 |
| 22 Feb. | 4 | 3 | 7 | -15.95 | 118.11 | 30 | 34.51 | 29.22 | 193.1 | 0.125 | 42 |
| 25 Feb. | St 37-Argo4 | na | 8 | -15.01 | 117.44 | 2 | 34.59 | 30.39 | 186.7 | 0.050 | 42 |
| 28 Feb. | St 44-Argo11 | na | 9 | -13.53 | 116.60 | 2 | 34.42 | 29.94 | 103.3 | 0.044 | 50 |

**Table 2.** General details of the grazing experiments (Exp). Duration of each incubation phase, mean number of copepods incubated per bottle, copepod weight, abundance of ^15^N-*Pro* (*Prochlorococcus*) and ^15^N-*Syn* (*Synechococcus*) inoculated in copepod bottles. Errors are standard deviations (sd).

| Cycle /<br>Station | Exp | Microzoo<br>incubation | Microzoo + copepod<br>incubation | Total<br>incubation | Copepods |  | <sup>15</sup>N-Enrichment |  |
| --- | --- | --- | --- | --- | --- | --- | --- | --- |
|  |  | T<sub>0</sub>-T<sub>24</sub> | T<sub>24</sub>-T<sub>36</sub> | T<sub>0</sub>-T<sub>36</sub> | Abundance | Weight | <sup>15</sup>N-*Pro* | <sup>15</sup>N-*Syn* |
|  |  | (h) |  |  | (cop. bottle<sup>-1</sup>) | (µg C ind<sup>-1</sup>) | (cells mL<sup>-1</sup>) |  |
| 1 | 1 | 23.0 | 12.2 | 35.2 | 48 ± 5 | 24.1 ± 7.5 | 13,400 |  |
| 2 | 2 | 21.4 | 6.0 | 27.4 | 38 ± 5 | 8.4 ± 0.6 | 6,730 |  |
| 2 | 3 | 21.5 | 14.6 | 36.1 | 70 ± 0 | 34.1 ± 12.8 | 6,710 |  |
| 3 | 4 | 22.7 | 13.5 | 36.2 | 61 ± 4 | 56.0 ± 19.3 | 5,680 | 25,680 |
| 3 | 5 | 23.8 | 12.9 | 36.7 | 39 ± 4 | 24.2 ± 8.6 | 5,660 | 57,090 |
| 4 | 6 | 22.4 | 13.5 | 35.9 | 26 ± 2 | 17.2 ± 1.5 | 8,540 | 73,800 |
| 4 | 7 | 22.5 | 14.2 | 36.7 | 32 ± 5 | 26.2 ± 5.2 | 9,480 | 66,600 |
| 37 | 8 | 21.1 | 12.8 | 33.9 | 11 ± 1 | 20.6 ± 9.3 | 10,970 | 89,940 |
| 44 | 9 | 21.7 | 13.1 | 34.8 | 27 ± 9 | 104 ± 18 | 10,940 | 91,160 |

**Figure 1.**
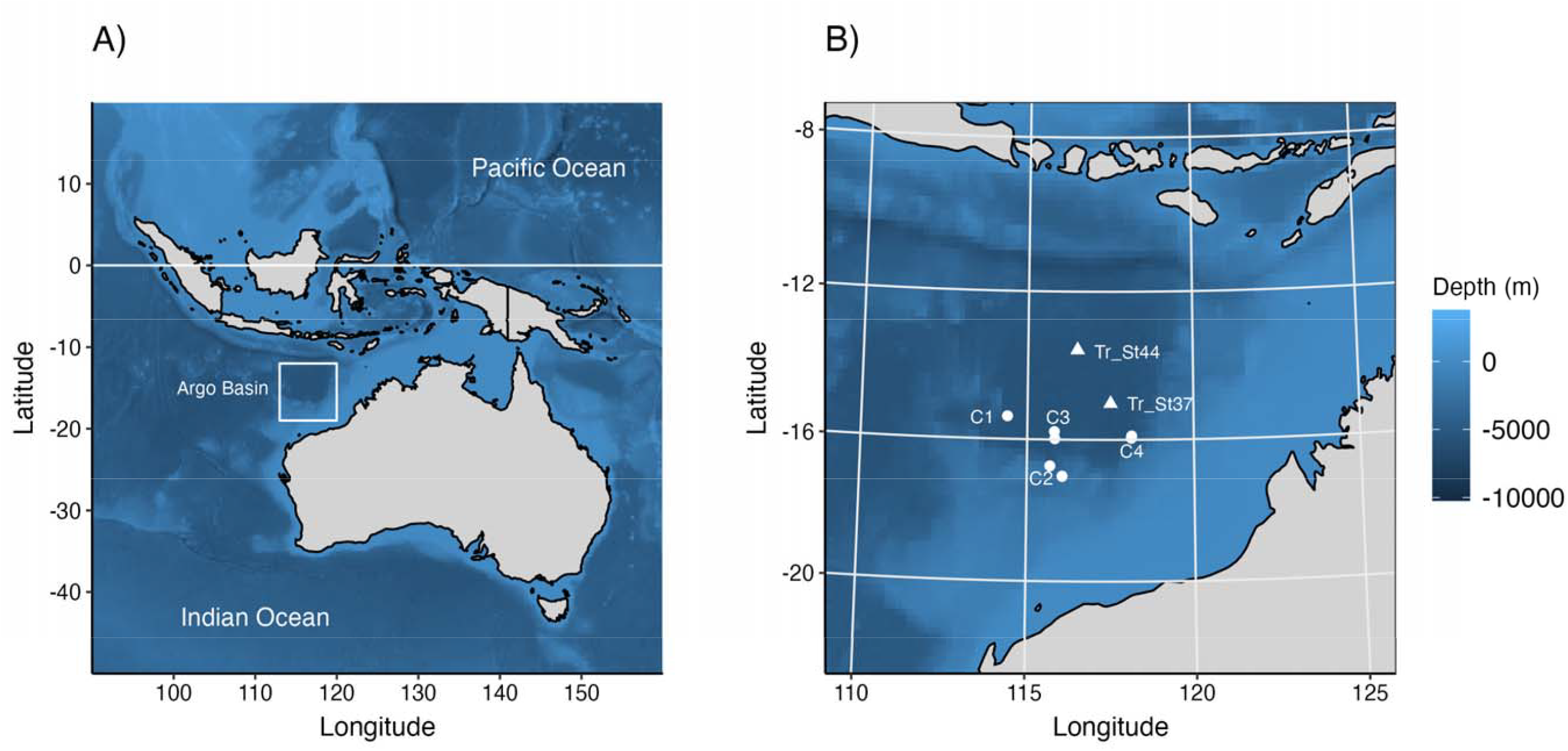
A) BLOOFINZ research cruise study area. B) Geographical position where grazing experiments were conducted, which corresponds to the region enclosed within the square in A). Circles indicate cycle experiments; triangles indicate transit experiments.

**Figure 2.**
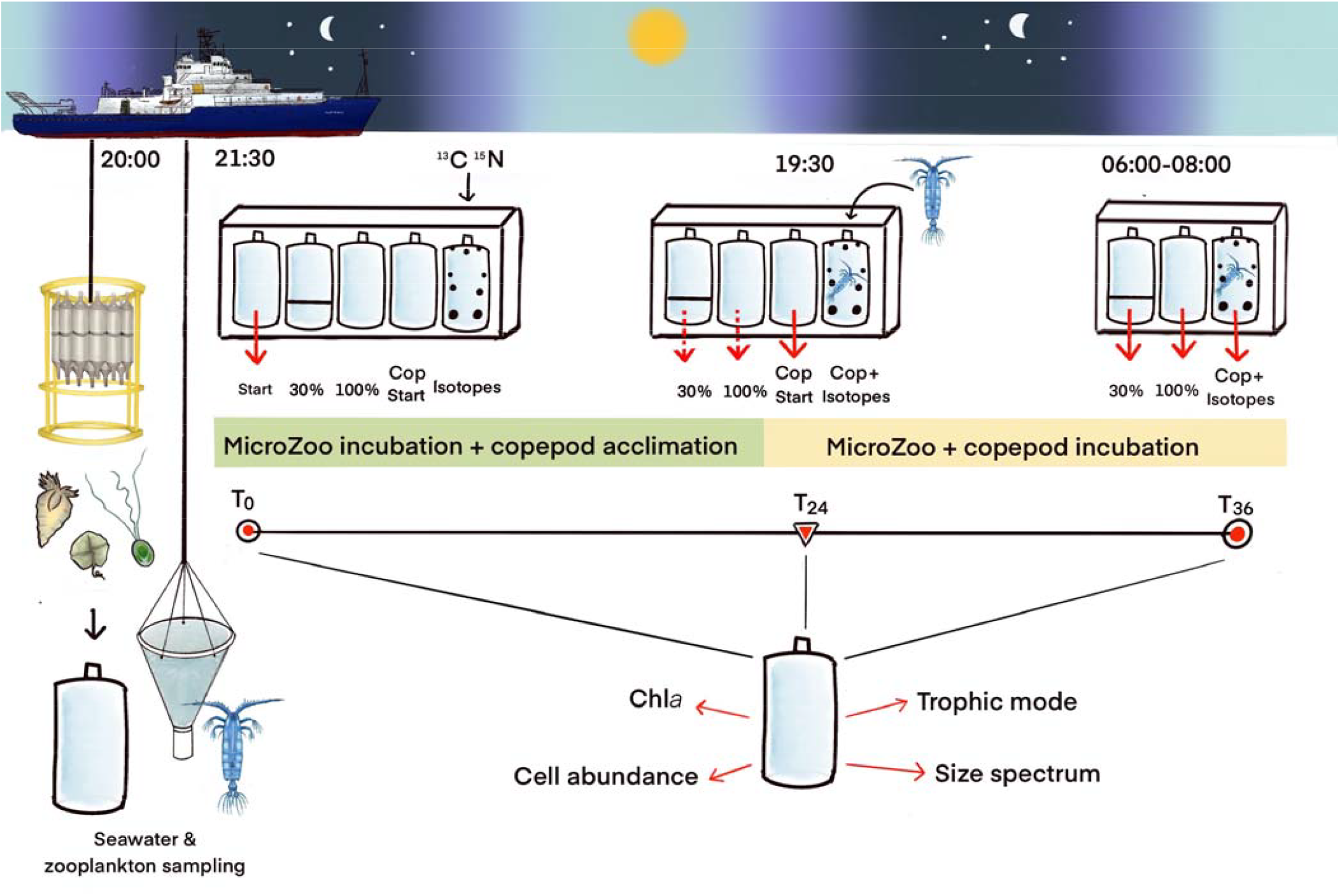
Timeline of the experimental phases. Evening sampling of seawater and zooplankton, filling of bottles and isotope spike at the onset of the Microzoo incubation (Start, T_0_); 24h later, acclimated zooplankton were then added in the isotope-containing bottles, and start of the Microzoo+copepod incubation (T_24_). After 12h incubation with copepods, termination of the experiment (End, T_36_). Red arrows indicate the bottles sampled for analyses. The illustration is merely a representative scheme, since triplicate bottles per treatment were used.

### 2.2. Seawater and zooplankton sampling

Seawater for grazing incubations was typically collected from Niskin bottles on CTD casts in early evening (19:00 to 20:30 local; Fig. 2). During cycles (Exps 1-7), low-Chl*a* low-nutrient water was sampled between 28 and 30 m depth. For the two transit experiments (Exps 8-9), water was collected from the surface (2 m depth) using trace metal-clean bottles (Table 1).

Between 20:30 and 23:00, after seawater collection, zooplankton live samples were collected (Fig. 2) with gentle vertical tows (∼10 m min^-1^) using a 1-m^2^ ring net (200-µm mesh), equipped with a non-filtering 30-L cod end and depth sensor attached to the net frame. Sampling depths ranged from 38 to 60 m depending on the experiment (Table 1). The animals were carefully placed in two 20-L buckets, diluting them by 1/3 with fresh filtered seawater, and allowed to acclimate for ∼22 h in a temperature-controlled room (set to the average sea surface temperature) in the darkness until sorted for experiments.

### 2.3. Bottle treatments and experimental procedures

Each experiment included: a) three initial “Start” bottles, filled with seawater and immediately sampled after filling (T_0_); b) three “30%” bottles containing 30% seawater and 70% 0.1 µm-Acropack filtered seawater (FSW); c) three “100%” bottles with undiluted seawater; d) three “CopStart” bottles including undiluted seawater, representing the starting conditions for copepod incubations; e) three “Cop+Isotopes” bottles containing undiluted seawater, ^15^N (nitrogen)-labelled prey and ^13^C (carbon) bicarbonate inoculated at T_0_, and copepods, which were added ∼22-24 h later (at T_24_; Fig. 2). The use of dual isotopic tracers was meant to identify the source of copepod nutrition i.e., autotrophic diet associated with ^13^C signal while phagotrophic nutrition (transfer from cyanobacteria to copepods through intermediate consumers) linked to ^15^N signal. We assumed that copepods were not able to feed on cyanobacteria directly due to their suboptimal prey size.

#### Microzoo incubation (T_0_-T_24_)

Water from the evening CTD cast was transferred with acid-washed silicone tubes into 2.7-L acid-washed transparent polycarbonate bottles following three seawater rinses. The bottles were filled to capacity avoiding bubbles and ensuring a slow and gentle flow. For Exps 8 and 9, the water was first collected with trace metal-clean 5-L GO-Flo bottles, then gently homogenized in carboys and added to the 2.7-L bottles as for other experiments. During these two last experiments, 5 mL water was removed from all the bottles and replaced with 5 mL of underway surface seawater to ensure non-limiting trace metal concentration for photosynthesis.

Before the start of the cruise, ^15^N-labelled *Prochlorococcus* (*Pro*) and ^15^N-labelled *Synechococcu*s (*Syn*) were prepared in the laboratory by filtering ^15^N-labelled cell culture stocks onto 25-mm diameter 0.2-µm polycarbonate filters (ca. 2-8 × 10^8^ cells filter^-1^). Filters were flash-frozen in liquid nitrogen in 2-mL cryo-Eppendorf tubes and stored long-term at - 80°C. Prior to each experiment, the ^15^N-labelled cells from one filter were resuspended in 13-14 mL of 0.1-µm Acropack FSW, mixed by vortex, and refrigerated for ∼1 h before spiking (2-4 mL) experimental bottles to start (T_0_) the Microzoo incubation into the Cop+Isotope bottles (where copepods will be added on the next day; Fig. 2). Depending on the experiment, the type of cyanobacteria (*Pro* or *Syn*), the availability of the ^15^N-prey filters, and the ^15^N-labelled cyanobacteria recovered, prey concentrations that were added ranged from 6,700 to 102,000 cells mL^-1^ (Table 2). Additionally, 1 mL of ^13^C bicarbonate stock was inoculated to reach a final concentration of 0.25 mM ^13^C in the same Cop+Isotope bottles. For Exps 1-3, only ^15^N-*Pro* was added, whereas Exps 4-9 also received ^15^N-*Syn*, usually in concentrations that were 5 to 10-fold those of ^15^N-*Pro* (Table 2). After filling and adding labelled-prey, the bottles were placed in a deckboard incubator exposed to 25% of incident PAR to match the irradiance at the depth of seawater collection. The incubators were covered at night to ensure complete darkness, and three Start bottles were sampled for initial conditions of the different abiotic and biotic parameters.

#### Microzoo+copepod incubation (T_24_-T_36_)

Copepods were sorted out of the cod end community after ∼20-24 h of conditioning in a constant-temperature room, by concentrating the sample onto 200-µm mesh filters and transferring into small plastic jars containing seawater. Aliquots of the concentrated zooplankton sample were poured into Petri dishes and only healthy intact individuals were sorted with a glass Pasteur pipette under a dissecting microscope. The selected copepod specimens were calanoids (*Candacia* spp., *Euchaeta* spp., *Labidocera* spp., other calanoids) and cyclopoids (*Oncaea* spp., *Sapphirina* spp., and Corycaeidae). Sorted individuals were temporarily pipetted into 50-mL Falcon tubes prefilled with 40 mL of FSW and covered with aluminum foil prior to inoculation. The number of copepods incubated varied by experiment depending on availability (10-70; Table 2). Prior to introducing copepods into the experimental Cop+Isotope bottles, spiked with ^13^C and ^15^N-labelled prey the previous day, the sorted animals were concentrated down to 15-20 mL by 50 µm-reverse filtration, and 15-20 mL of the bottled water was removed with a serological pipette to allow for the addition of the copepods. The seawater volume containing the copepods was then gently mixed and transferred to the corresponding bottles. From the moment copepods were inoculated, the incubations lasted generally 12 h, except during Exp2 (6 h). Three Cop Start bottles were sampled for initial conditions, and the samples were processed according to the same protocols for the Microzoo incubation samples. An aliquot of copepods from the mixed community was stored on 50-µm mesh filters and frozen at -80ºC until analysis of their elemental contents and natural C and N isotope values (see Section 2.4). After ∼12 h from copepod addition (T_36_), all the bottles were removed from the deck incubator, quickly transported into the laboratory in the dark, and processed for final sampling following the same order in which they were previously filled and sampled.

### 2.4. Sample processing

At T_0_ and T_24_, experimental bottles were gently turned to keep seawater homogenized, and aliquots were collected for Chl*a* fluorescence and flow cytometric measurements of cell abundances (Fig. 2). At the end of copepod incubations (T_36_), the copepods were removed from the Cop+Isotope bottles by reverse filtration through 200-µm mesh, counted under a dissecting microscope, and filtered onto pre-weighed 50-µm sieves. Each filter was stored in a capped petri dish and immediately frozen (-80°C). The remaining water in the bottles was gently homogenized and subsampled as for the Microzoo incubation.

For Chl*a* estimates, 285 mL subsamples were collected in dark amber bottles and filtered onto 25-mm diameter GF/F Whatman filters. The filters were folded and the Chl*a* extracted in 5 mL 90% acetone kept in glass screw-cap tubes in a -20°C freezer in total darkness. After 24 h, the samples were vortexed, filters removed and Chl*a* fluorescence measured with a Turner Designs model 10AU Field fluorometer calibrated against a pure Chl*a* standard (Strickland and Parsons, 1972).

Flow cytometry (930 µL) samples were collected to estimate cell abundances of prokaryotes and protistan eukaryotes. Live runs were done to enumerate pico- and nanoplankton eukaryotes in the categories of picoautotrophs, nanoautotrophs, picomixotrophs, nanomixotrophs, and nanoheterotrophs. The live samples were stained with two probes: Hoechst 34580 (DNA-specific; Selph, 2021) and LysoTracker Green (LTG, acidic organelle-specific). The latter was used to detect the presence of food vacuoles in heterotrophic (Rose et al., 2004) and mixotrophic protists (Sato and Hashihama, 2019) on living cells since LTG is not retained in dead cells’ vacuoles (more details in Selph et al., this issue, a). After 5-10 min incubation with stains under dim light, the live samples were analyzed for particle concentration based on detectors for blue fluorescence from DNA (450±45 nm, excited with a 375 nm laser), green and red fluorescence of LTG (525±40 nm) and Chl*a* (690±50 nm), respectively, from a 488 nm laser, and forward and 90° side light scatter (FSC and SSC, respectively) using a portable Beckman Coulter CytoFlex S flow cytometer.

The distinction between eukaryotic populations was performed using Chl*a* fluorescence, light scatter, and signals from DNA and LTG staining (details in Selph et al., this issue, a). Specifically, autotrophs revealed positive Chl*a* signals and negative LTG signals (determined in comparison to maximum LTG signals from *Pro* and heterotrophic bacteria (Hbact) as negative controls), mixotrophs showed positive signals for both Chl*a* and LTG, and heterotrophs displayed negative Chl*a* signals and positive LTG signals. Eukaryotic size estimates were from FSC signals calibrated with various sizes of fluorescent polystyrene beads (0.5-10 µm diameter), with picoplankton defined as <2 µm and nanoplankton as >2 µm-20 µm.

For prokaryotes, an additional aliquot (930 µL) was preserved with 30 µL of 16% paraformaldehyde and immediately frozen at -80°C for a posteriori enumeration of Hbact, *Pro*, and *Syn*. These samples were thawed in batches on the ship and abundances determined by Hoechst 34580 DNA after staining 1 h before analyses (Selph, 2021).

Copepods, both initial and those in the grazing experiments, were dried and packaged in tin cups for elemental analyzer isotope-ratio mass spectrometry. Initials copepods were measured for natural δ^13^C and δ^15^N, while grazing experiment copepods were measured for enriched δ^13^C and δ^15^N. Depending on copepod size, we packaged between 2-8 copepods per tin cups to ensure sufficient material for analysis. To estimate the isotopic content of copepod food, particulate organic matter (POM) from the final spiked incubations was filtered onto pre-combusted GF/F filters (1100-1880 mL), dried at 60°C for 24 h, packed into tin cups, and sent for analysis. All isotopic samples were processed at the University of California (Davis).

### 2.5. Determination of microzooplankton grazing rates

The net growth rates (Eq. 1a, b) of prey in the 30% diluted samples (*k*_*d*_) and in the 100% undiluted samples (*k*) were calculated as follows:

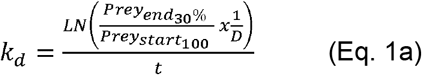

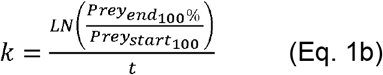

where *Prey*_*end*_ *and Prey*_*start*_ are prey concentrations (cells mL^-1^) at the end and start of the incubation respectively, *D* is the dilution factor, and *t* is the incubation duration (d^-1^).

Prey mortality by microzooplankton grazing rates (*m*, Eq. 2) were determined for Chl*a* and unicellular prey groups including prokaryotes (Hbact, *Pro, Syn*), and eukaryotes (picoautotrophs, nanoautotrophs, picomixotrophs, nanomixotrophs, nanoheterotrophs) using equations from Landry et al. (2008, 2025) as:

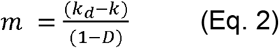

### 2.6. Determination of copepod feeding rates and food selection

Copepod grazing, clearance, and ingestion rates were calculated as in Frost (1972) based on prey disappearance relative to controls and later corrected for microzooplankton grazing following Nejstgaard et al. (2001). Since nano-microzooplankton were also actively grazing on the same prey community, prey availability for copepods was reduced compared to the initial standing stock. To account for this and to avoid underestimating copepod grazing, we additionally applied the correction proposed by Nejstgaard et al. (2001) (Eq. 3, Eq. 4). Nanoheterotrophs and nanomixotrophs (total nanophagotrophs) were assumed to exert the main grazing impact within the microzooplankton community analyzed, not only for their abundance but also for their phagotrophic potential over picograzers:

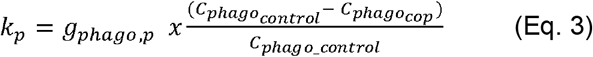

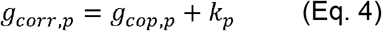

where *g*_*phago,p*_ is the grazin g by nanophagotrophs on each prey type, *C*_*phago*_ is the mean nanophagotroph concentration in the control and in the copepod bottles, and *g*_*cop,p*_ is the grazing by copepods on each prey type. Finally corrected clearance rate (*F*, mL ind^-1^ d^-1^; Eq. 5) and ingestion rate (*I*, cells ind^-1^ d^-1^; Eq. 6) were calculated according to Frost (1972) as:

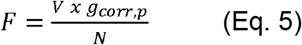

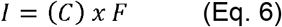

where *V* is the total volume of the incubation (mL), *N* is the number of live copepods retrieved after the incubation and *(C)* is the average prey concentration (cells mL^-1^) measured between start and end of the incubation in the copepod grazing bottles.

Food selection by copepods was expressed through the Chesson’s α index (Chesson, 1978) to determine preference for the five eukaryotic prey groups and through the Ivlev’s *E* index (Ivlev, 1961) for trophic and size classes separately. Both α and *E* and were calculated according to equations 7 and 8, respectively.

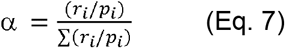

where Σ(*r*_*j*_/*p*_*j*_) is the sum of all prey types’ relative availability. α ranges from 0 to 1, with equal preference being α =1/n (where n is the number of prey types, in our case, α = 1/5 = 0.2).

Values of α>0.2 indicate preference for prey type *i*, whereas α<0.2 represents avoidance of prey type *i*.

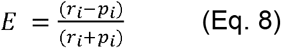

where *r*_*i*_ is the proportion of prey type *i* in the predator’s diet, and *p*_*i*_ is the proportion of prey type *i* in the environment. *E* ranges from -1 to +1, being *E*>0 indicative of preference for prey type *i*, and *E*<0 indicative of prey *i* avoidance, whereas *E*=0 indicates neutral ingestion, which corresponds to equal preference. Both indices give a measure of how much an ingested prey is preferred or avoided by copepods in proportion to its relative contribution in the environment. However, Chesson’s α index takes rare groups of prey into account, whereas Ivlev’s *E* index does not. Prokaryotes (Hbact, *Pro, Syn*) were excluded from the copepod grazing analyses due to negligible or null consumption.

Prey abundance and copepod ingestion rates were converted to carbon (C) biomass units from literature equations. Prokaryote conversions were: Hbact: 5 fg C/cell (Gundersen et al., 2002), *Pro*: 32 fg C/cell (Garrison et al., 2000), and *Syn*: 101 fg C/cell (Garrison et al., 2000). Eukaryotic conversions used the mean equivalent spherical diameter (ESD, µm) measured by epifluorescence microscopy to estimate biovolume (Yingling et al., this issue), then converted to carbon biomass using the Menden-Deuer and Lessard (2000) relationship in each cycle for the different groups of eukaryotes, namely picoeukaryotes: 0.32 pg C/cell (mean biovolume of 0.8-2 µm cells), auto- and mixotrophic nanoeukaryotes: between 5 and 9 pg C cell^-1^, and heterotrophic nanoeukaryotes: 5 pg C cell^-1^. A carbon conversion of 5 pg C cell^-1^ was used for heterotrophic nanoeukaryotes based on mean 3.74-µm ESD of these cells (Yingling et al., this issue). For transit experiments (Exps 8 and 9), no data were available from epifluorescence samples; hence, cell C contents for nanoauto- and nanomixotrophs were inferred from FChl*a*:C ratios measured at the surface during cycles and calculated according in proportion to those measured during transits (6 pg C cell^-1^).

### 2.7. Grazing impact by microzooplankton and copepods

The percent contributions of microzooplankton to the daily removal of unicellular planktonic production were calculated as:

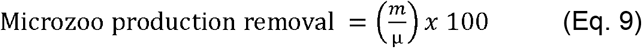

where *m* is the grazing rate (d^-1^) by microzooplankton on a specific prey item and *µ* is the growth rate (d^-1^) of the prey type (Landry et al., 2025), calculated as:

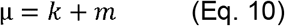

The percent contributions of copepods to the daily removal of protist production were calculated using clearance rate (*F*, L ind^-1^ d^-1^), copepod abundance in our incubations (*N*, ind L^-1^), the abundance of each prey (*Cp*, cells L^-1^), and total prey (*TotC*, cells L^-1^):

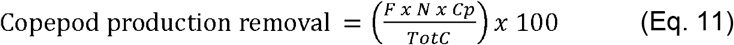

### 2.8. Copepod consumption of isotopically labelled prey

To obtain independent estimates of copepod trophic reliance on auto- vs mixo-heterotrophic diet, we estimated the pathway of C ingestion from primary producers and the consumption of N grazers of ^15^N-labelled *Pro* and *Syn*. The copepod biomass (*W*, µg ind^-1^) and isotopic signals for δ^13^C and δ^15^N of both prey (*food_at*, %) and copepods (*cop_at*, %) were provided directly from the isotope-ratio mass spectrometry analyses. The values used for ‘*food_at*_*T36*_’ were the POM values taken from the copepod incubation bottles at the end of the experiment.

The daily copepod C and N ingestion rates (*IR*, µg ind^-1^ d^-1^, Eq. 12) were calculated as the difference in copepod δ^13^C and δ^15^N isotopic signal between the start (T_24_) and the end (T_36_) of the incubation (*t*, hours) and appropriately subtracting the corresponding baseline food isotopic signal from the copepod isotopic signal (Verschoor et al., 2005). The percentage of ingested biomass to the total copepod biomass gives the C- and N-specific daily ration (*DR*, % d^-1^, Eq. 13).

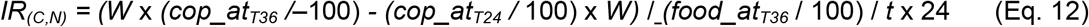

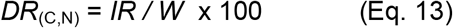

### 2.9. Data analysis and statistics

Data were checked for normality (Shapiro-Wilk’s test) and homoscedasticity (Bartlett’s test) assumptions. Differences in copepod feeding rates (clearance, ingestion) and selectivity and microzooplankton grazing rates were tested as a function of food quality indicators: prey trophic mode and size class. One-Way ANOVA and/or non-parametric Kruskal-Wallis tests were applied to test a priori differences in variables between trophic modes (autotrophic-mixotrophic-heterotrophic) while *t*-tests were used to compare size classes (picoplankton-nanoplankton). For tests yielding significantly different variance between trophic groups, post-hoc Tukey HSD and/or Dunn’s tests were applied for multiple comparisons. Differences were considered significant at the 0.05 level. Graphical exploration of results and statistical analyses were performed using RStudio v4.2 (R Core Team, 2021).

## 3. Results

### 3.1. Food landscape: biomass, size and trophic spectra of unicellular plankton

At the start of the microzooplankton incubation (T_0_), Chl*a* biomass ranged from 0.05 to 0.1 µg L^-1^, with slightly lower concentrations for the shallower depths sampled for transit experiments (0.05-0.06 µg Chl*a* L^-1^) compared to cycle experiments (0.07-0.1 µg Chl*a* L^-1^) (Table3). The prey community available for microzooplankton was numerically dominated by Hbact (482,000 – 625,000 cells mL^-1^) and *Pro* (128,000-276,000 cells mL^-1^; Table 3). *Syn* (1,260-3,460 cells mL^-1^) had similar abundances as nanoheterotrophic eukaryotes (817-2610 cells mL^-1^). In contrast, the remaining eukaryotic groups represented by nanomixotrophs (28-666 cells mL^-1^), picomixotrophs (19-282 cells mL^-1^), picoautotrophs (29- 202 cells mL^-1^), and nanoautotrophs (28-169 cells mL^-1^) were much less abundant (Table3). On average, Hbact accounted for 70-80% of microzooplankton prey, followed by *Pro* (20-30%), whereas *Syn* and all the eukaryotic populations were negligible (<1%) in numerical terms.

**Table 3.**
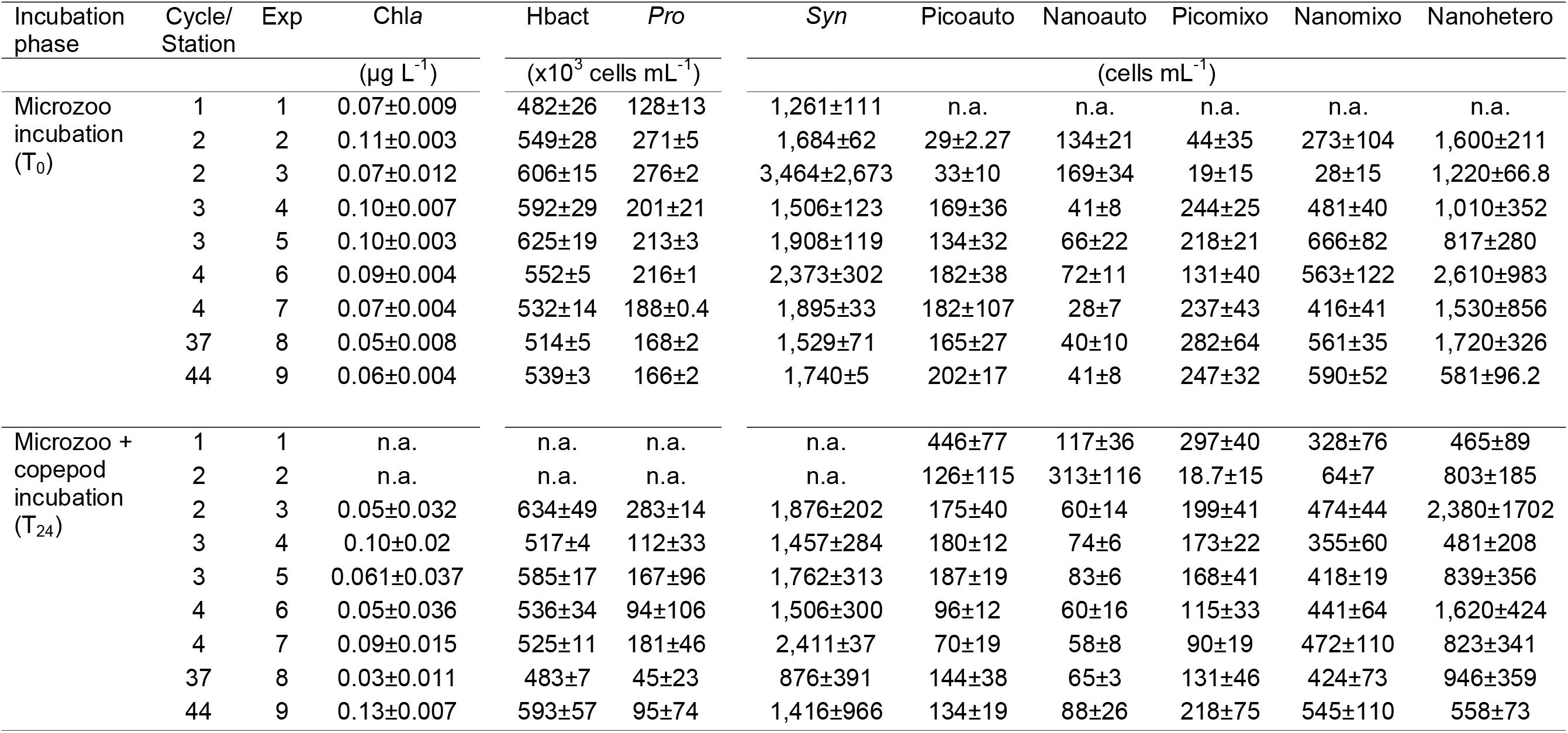
Concentration of Chl*a* (µg L^-1^) and abundance of plankton groups (cells mL^-1^) at the start of Microzoo incubation (T_0_) and at the start of Microzoo+copepod incubation (T_24_). Samples not analysed: n.a. Data are means ± sd (*N*=3). Abbreviations: Exp: experiment; Chl*a*: Chlorophyll *a*; Hbact: heterotrophic bacteria; *Pro: Prochlorococcus; Syn: Synechococcus*; Picoauto: picoautotrophs; Nanoauto: nanoautotrophs; Picomixo: picomixotrophs; Nanomixo: nanomixotrophs; Nanohetero: nanoheterotrophs.

In terms of carbon (C), unicellular community biomass was higher in Cycle 2 (21 µg C L^-1^) and Cycle 4 (24.5 µg C L^-1^) compared to Cycle 3 and transit experiments (∼17.5 µg C L^-1^) Fig. 3A). *Pro* and nanoheterotrophs contributed most (30-40% each), nanomixotrophs and Hbact were slightly lower (20-30% each), and *Syn*, picoautotrophs, picomixotrophs and nanoautotrophs together comprised <10% of total biomass. Overall, unicellular heterotrophs accounted for 58% of the C standing stock, whereas autotrophs were 18 to 32% and the remaining 8 to 22% were mixotrophs (Fig. 3B). Prokaryotes and smaller eukaryotes (picoplankon) were the most abundant but contributed less to total biomass (25-40%) than nanoplankton (58-72%) (Fig. 3C).

**Figure 3.**
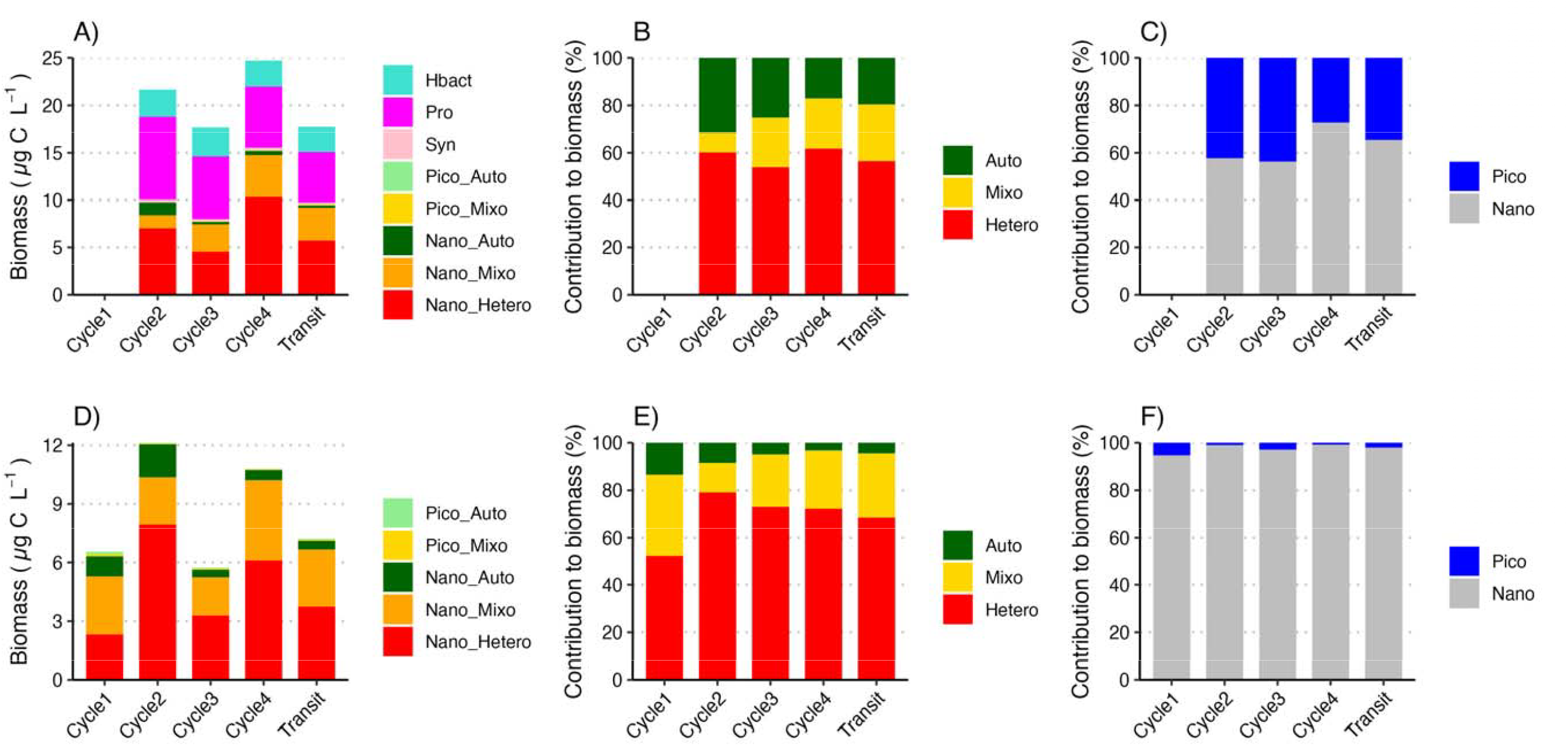
Initial biomass of the unicellular groups of pico- and nanoplankton available as prey for microzooplankton at T_0_ (A,B,C) and copepods at T_24_ (D,E,F) and relative contribution of trophic modes and size classes to the carbon biomass in cycle and transit experiments. Data are means (*N*=3-6). Note that during Cycle1, data on eukaryotes abundance at T_0_ were not available.

At the start of the copepod incubation (T_24_), Chl*a* biomass varied between 0.03 and 0.13 µg Chl*a* L^-1^ (Table 3). Concerning the prey populations, eukaryotic groups accounted for a total biomass of 5.6-12 µg C L^-1^, higher in Cycle 2 (12 µg C L^-1^) and Cycle 4 (10.6 µg C L^-1^) compared to Cycle 1, Cycle 3, and transit experiments (5.8-7.3 µg C L^-1^), in line with initial biomass (T_0_; Fig. 3D). Heterotrophic protists accounted for 53-79% of eukaryotic biomass, followed by mixotrophs (12-33%), and autotrophs (3-15%) (Fig. 3E). The prey biomass available to copepods, i.e. sampled in bottles 24 h later, was mostly nanoplankton (93-99%) rather than picoplankton (1-7%) (Fig. 3F).

### 3.2. Copepod feeding on eukaryotes: ingestion, diet composition and selectivity

After the copepod incubation, Chl*a* biomass remained stable in control bottles and increased by ∼30-60% in copepod grazing bottles (not shown); hence, grazing of copepods based on Chl*a* disappearance was not estimated. Total grazing on eukaryotes oscillated between 0.43 and 1.64 d^-1^ (median= 1.31±0.23 d^-1^; Fig. 4A), with clearance rates ranging from 18.5 to 336 mL ind^-1^ d^-1^ (median= 80.7±55.7 mL ind^-1^ d^-1^; Fig. 4B). Significant differences in copepod feeding emerged in both grazing (*m*) and clearance (*F*) between hetero-and mixotrophs (*m*: *p*=0.0058; *F*: *p*=0.0045) and between pico- and nano sized prey (*m*:, *p*<0.0001; *F*: *p*=0.0029). Considering all prey types, between 3.4 and 138 ng C ind^-1^ d^-1^ (median= 31.4±16 ng C ind^-1^ d^-1^) were ingested by copepods in total including nanomixotrophs (0-12417 cells ind^-1^ d^-1^, 0-74.5 ng C ind^-1^ d^-1^) and nanoheterotrophs (0-5,300 cells ind^-1^ d^-1^, 0-26.5 ng C ind^-1^ d^-1^) at higher rates compared to picomixotrophs (356-12,400 cells ind^-1^ d^-1^, 0.11-3.96 ng C ind^-1^ d^-1^), picoautotrophs (390-30,200 cells ind^-1^ d^-1^, 0.12-9.68 ng C ind^-1^ d^-1^) and nanoautotrophs (0-339 cells ind^-1^ d^-1^, 0-3.05 ng C ind^-1^ d^-1^) (Fig. 4C,D). Cell-specific ingestion increased linearly with prey abundance (*p*<0.001, *R*^2^=0.25, not shown), and differed significantly as functions of trophic mode (hetero-mixo, *p*=0.017) and size class (*p*=0.006). C ingestion scaled linearly with prey biomass (*p*<0.001, *R*^2^=0.49, not shown) and according to prey size (*p*=0.021).

**Figure 4.**
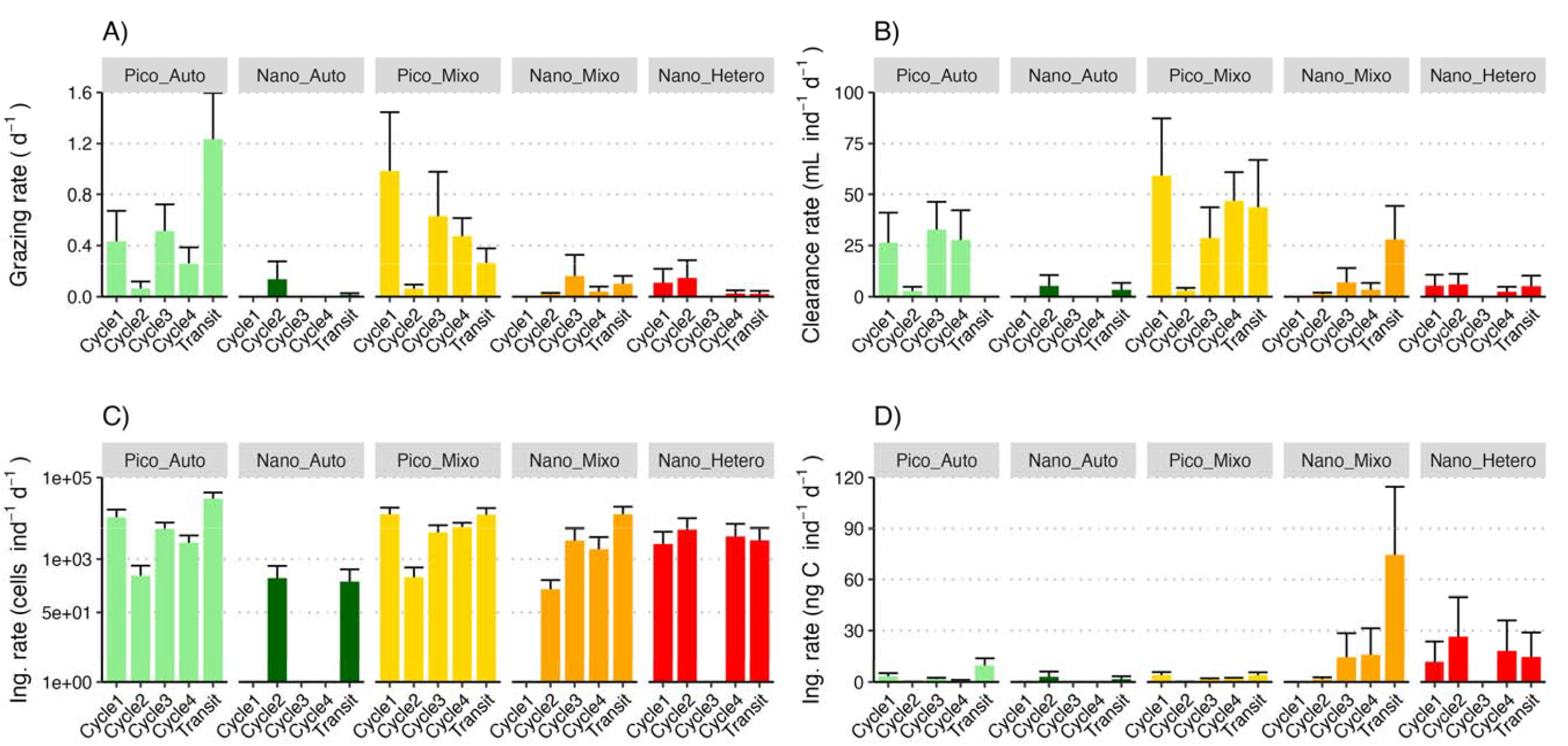
Copepod feeding. A) Grazing rate, B) clearance rate, C) cell-specific ingestion rates and D) carbon-specific ingestion rate on the different protist populations in cycle and transits experiments. Absent values indicate null or negative grazing. Data are mean ± standard error (*N*=3-6).

Copepods consumed a diverse diet, including a considerable proportion of each group in cell abundance terms (Fig 5A). Numerically, picoplankton cells contributed to a higher proportion of diet than nanoplankton (Fig. 5E), but nanoplankton dominated C intake (61.7-99.2%) (Fig. 5F). Among nanoplankton categories, C consumption derived mainly from nanoheterotrophs (0-84.3%) and nanomixotrophs (0-81.5%), leaving smaller fractions of C acquired from ingestion of picomixotrophs (0.4-20.8%), picoautotrophs (0.4-17.5%), and nanoautotrophs (0-9.7%) (Fig 5B). Higher contributions of nanoheterotrophs to copepod diet were recorded for Cycle 1,2 and 4 experiments, whereas an enhanced contribution of nanomixotrophs to copepod diet emerged in Cycle 3 and transit experiments. Despite the slight differences between cell-based and C-based estimates, mixotrophic (5.5-89.9%) and heterotrophic (0-84.3%) protists accounted for the highest contributions to copepod diets while strict autotrophs contributed substantially less (2.2-17.5%) (Fig. 5C,D).

**Figure 5.**
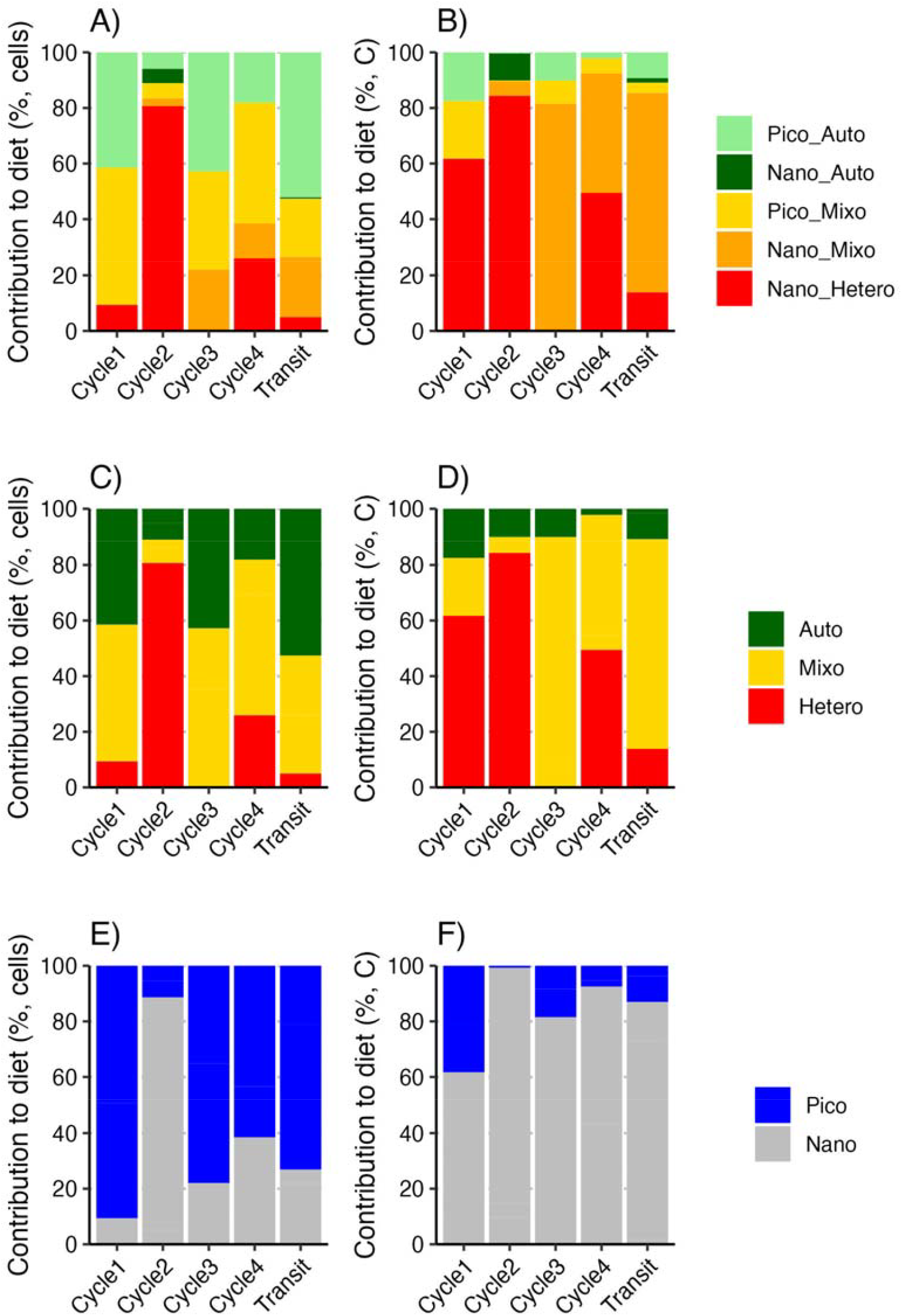
Copepod diet composition on eukaryotes in the different cycles and transit experiments. Data are mean relative contributions of groups (A, B), trophic modes (C, D), and size classes (E, F) to the copepod diet expressed as cell-specific ingestion (left panels) and carbon-specific ingestion (right panels).

Copepods showed neutral selection (α=∼0.2) for nanomixotrophs and picoautotrophs, while nanoheterotrophs, nanoautotrophs and picomixotrophs tended toward negative selection (α<0.2, Fig. 6A). Overall, however, no significant differences were found in selectivity among prey groups (*p*=0.894). Autotrophs were ingested in proportion to their availability (*E=+*0.01), heterotrophs were neglected (-1) and mixotrophs were moderately preferred (+0.31; Fig. 6B), but these differences among trophic categories were not significant due to high data variability (*p*=0.289). Positive moderate selection of copepods was shown for picoplankton (+0.3) at the expense of nanoplankton (-0.05), but this was not significant (*p*=0.563; Fig 6C).

**Figure 6.**
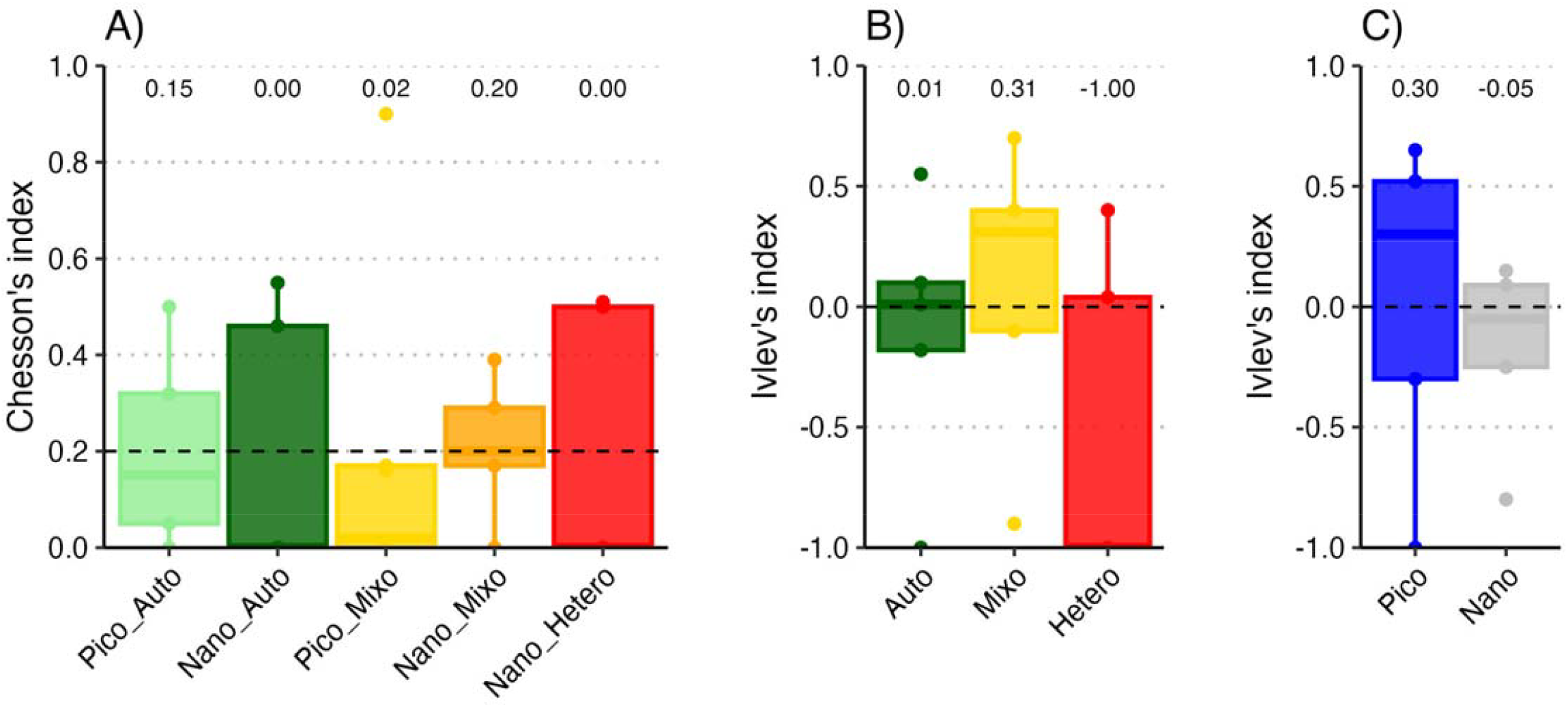
Copepod food selection. Chesson’s selectivity (α) index on A) the prey groups, and Ivlev’s electivity index (*E*) on the prey groups based on B) trophic mode, and C) size. Mean values from all experiments were pooled together per each category. Median values are reported for each prey and/or category. Isolated dots represent outliers. The dashed line represents neutral selection.

The daily total C ingestion rates of copepods based on bottle-incubation disappearance estimates ranged between 0.02 and 0.1 µg C ind^-1^ d^-1^ (equivalent to 20-100 ng C ind^-1^ d^-1^; Fig. 7A). The daily rations were generally very low, ranging between 0.05 and 0.34% d^-1^ (Fig. 7B). In contrast, estimating consumption of POM using ^13^C as a tracer yielded C-specific ingestion rates (Fig. 7C) and daily rations (Fig. 7D) that were one order of magnitude higher compared to those obtained using prey abundances analysed by flow cytometry. These ranged from 0.25 to 0.42 µg C ind^−1^ d^−1^ (250-420 ng C ind^-1^ d^-1^), which represented 0.8-2.4% d^-1^ of the copepod C biomass. Estimates of N ingestion through ^15^N-labelled prey ranged 0.01-0.07 µg N ind^−1^ d^−1^, (10-70 ng N ind^-1^ d^-1^) with higher values observed in Cycle 3, 4 and transit experiments (Fig. 7E) when the larger labelled prey (*Syn*) was used. N daily rations accounted for 0.25-1.0% d^-1^ (Fig. 7F). The daily ration for C (% body carbon) and N (% body nitrogen) based on the isotope tracers was ∼1% for both elements during Cycle 3, 4, and transits (Fig. 7D, F).

**Figure 7.**
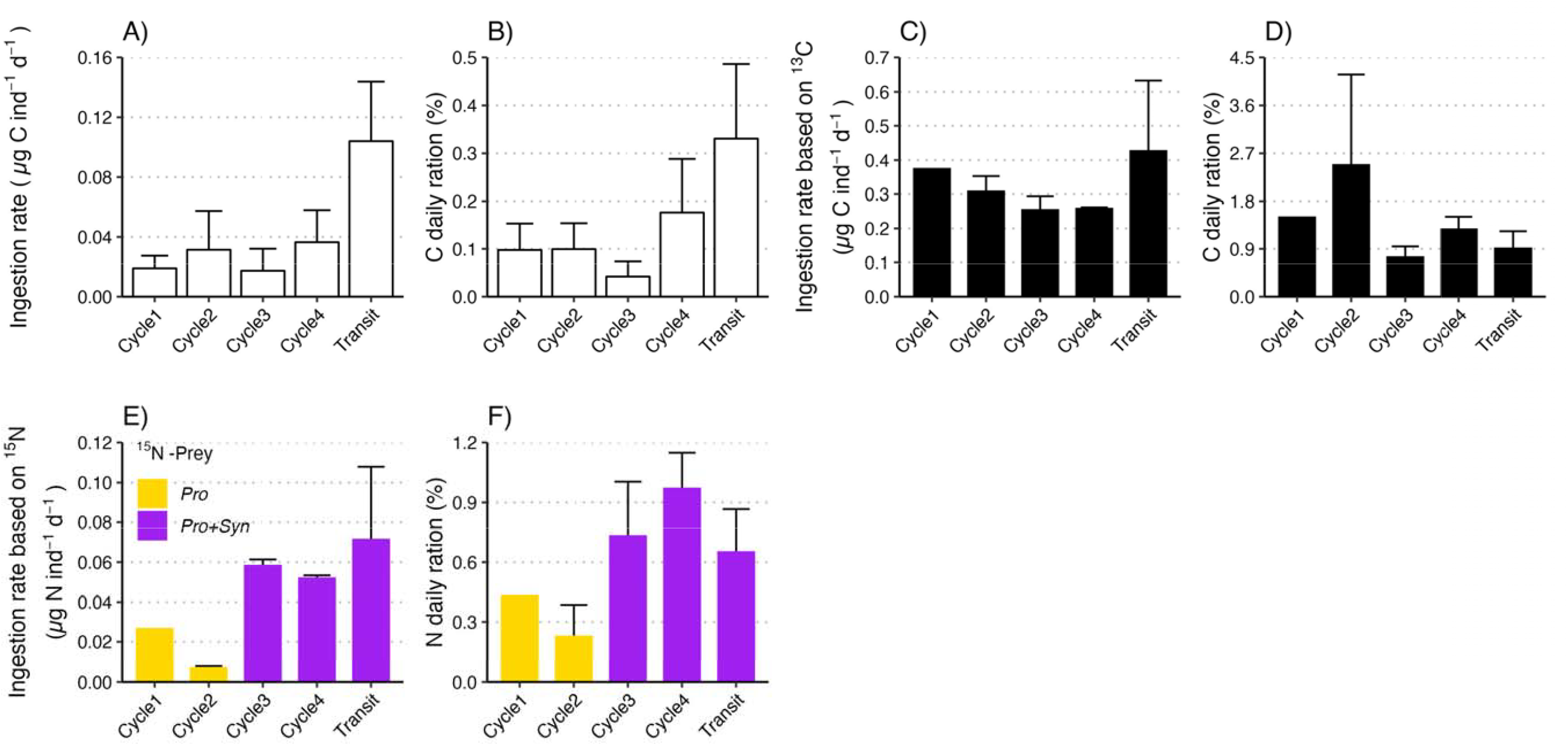
Copepod ingestion rate and daily ration. A) carbon-specific ingestion rate and B) daily ration estimated by food removal (black), C) carbon-specific ingestion rate and D) daily ration inferred through isotope signal (δ^13^C); E) nitrogen-specific ingestion rate, F) and daily ration inferred through isotope signal (δ^15^N). The concentration of ^15^N-*Pro* was negligible (10-13%) compared to that of ^15^N-*Syn* (87-90%) in Cycle 3, Cycle 4, and Transit. Data are mean±standard errors (*N*=9-18).

### 3.3. Microzooplankton grazing on prokaryotes and eukaryotes

Microzooplankton grazed variably on Chl*a* (0.34-0.66 d^-1^) with lower values in Cycles 1 and 4 compared to the other experiments but was not significantly different throughout the cruise (*p*=0.195; Fig. 8A). *Pro* (0.36-2.14 d^-1^), *Syn* (0.36-1.26 d^-1^) and Hbact (0.48-1.46 d^-1^; Fig. 8B) were grazed similarly by microzooplankton with some variability between experiments. Among eukaryotes, nanoheterotrophs were the most grazed prey (0.92-2.48 d^-1^), especially in Cycle 3 and along transits, followed by nanomixotrophs (0.48-1.24 d^-1^), nanoautotrophs (0.18-1.02 d^-1^; Fig. 8C), picoautotrophs (0.06-1.51 d^-1^) and picomixotrophs (0.08-0.93 d^-1^). Considering only eukaryotes, microzooplankton grazed on heterotrophs at significantly higher rates compared to autotrophs (*p*=0.0002), and mixotrophs (*p*<0.0001). Similarly, picoeukaryotes were grazed at significantly lower rates compared to nanoeukaryotes (*p*<0.0001). Considering both prokaryotes and eukaryotes, grazed biomass did not differ significantly between trophic modes (*p*=0.927), but it did between size classes (*p*=0.0002). The highest contribution to diet consisted mostly of nanoeukaryotes, namely nanoheterotrophs (24.7-58.6%), nanomixotrophs (13.8-40.1%) and nanoautotrophs (7.5-32.9%), whereas the other prokaryotic and eukaryotic prey accounted for <7% of the food intake (Fig. 8D). Out of the total grazed biomass, 24.8-58.7% was heterotrophic, 13.9-41.7% was mixotrophic, and a lower fraction (12.1-33.5%) originated from autotrophic nutrition (Fig. 8E). As the cruise progressed, increased heterotrophic relative dietary contributions were accompanied by a relative decreases in autotrophic diet, whereas the contribution of mixotrophs did not show a clear pattern. Nanoplankton contributed the highest microzooplankton food intake (95.1-97.9%) as opposed to picoplankton (2.1-4.9%) (Fig. 8F).

**Figure 8.**
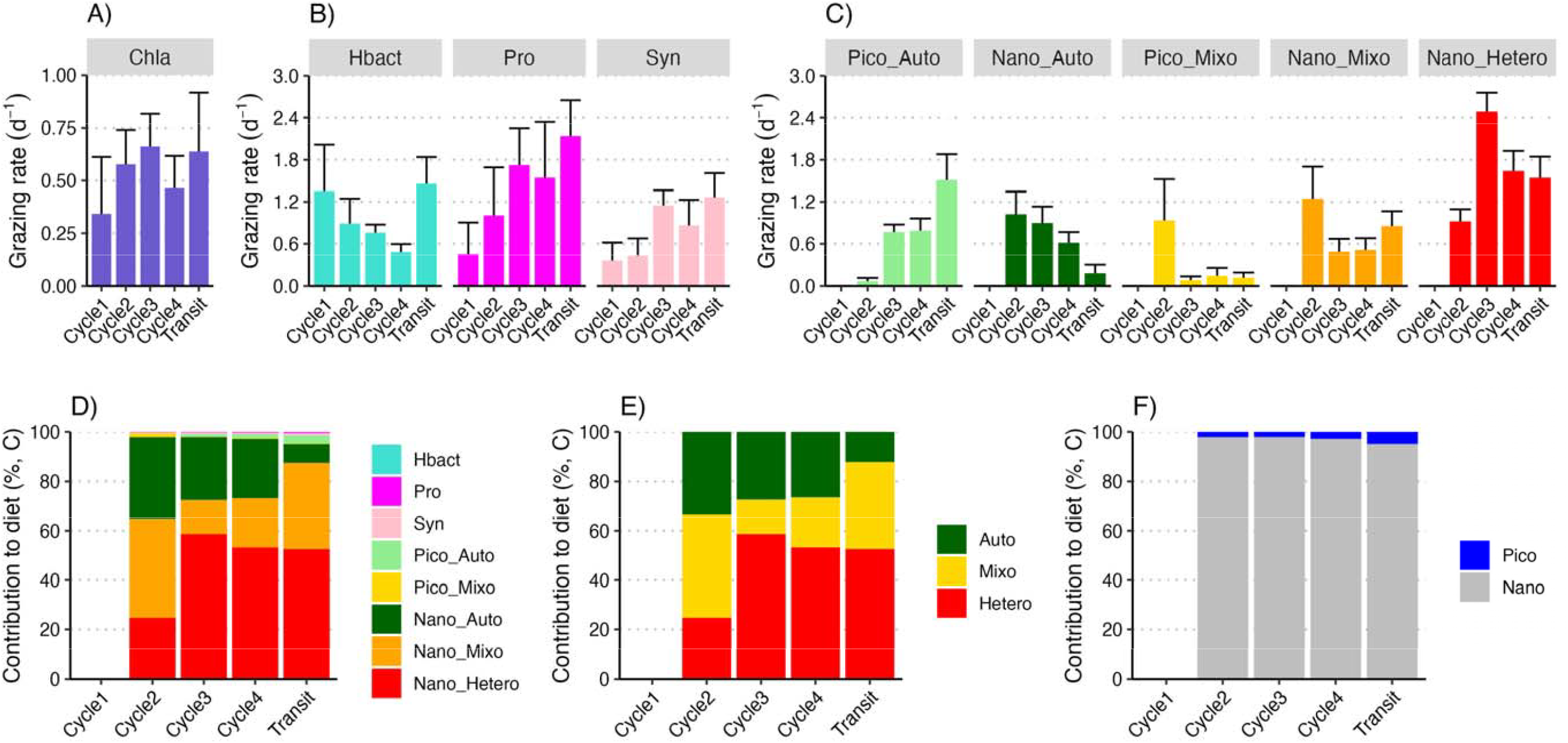
Grazing rate by microzooplankton on A) Chl*a*, B) prokaryotes, and C) eukaryotes and contribution to dietary carbon of the different D) prey groups, E) trophic modes and F) size classes. Data in the upper panels are mean ± standard error (*N*=3-6). Note that during Cycle1, data on eukaryotes abundance at T_0_ were not available, thus grazing rates were not measured.

### 3.4. Daily removal of unicellular plankton production

On average, 110% of the daily Chl*a* production was removed by microzooplankton (Fig. 9A), and 104%, 149% and 125% of the daily production of Hbact, *Pro* and *Syn* were consumed, respectively. Among eukaryotes, microzooplankton contributed to the high removal of nanoheterotrophs (149%), picoautotrophs (109%), nanoautotrophs (128%). Between 96% (pico) and 76% (nano) of daily mixotrophic production was removed by microzooplankton. By comparison, the highest copepod grazing impact on eukaryotes was exerted on picoautotrophs (13.8%), followed by picomixotrophs (9.6%), nanoheterotrophs (7.6%), nanomixotrophs (3.9%), while production of nanoautotrophs was the least impacted by copepod grazing (<0.1%; Fig. 9B).

**Figure 9.**
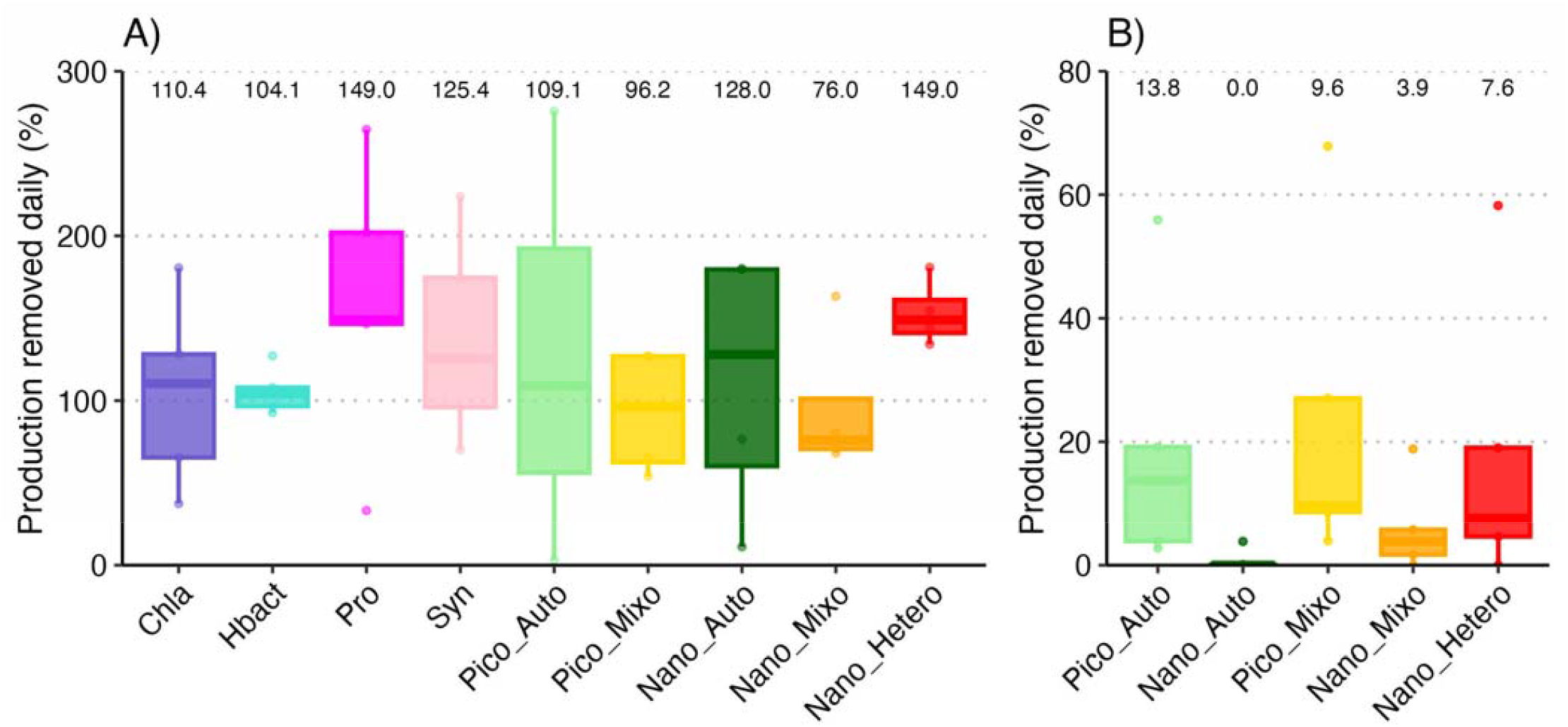
Percentage of prey production removed daily by A) microzooplankton over 24h, and B) copepods over the following 12h. Median values are reported on the top of each group in each station (*N*=5, including cycle and transit experiments).

## 4. Discussion

### 4.1. Community biomass, size classes and trophic modes

The Chl*a* and microbial standing stock estimates in our incubations align with values calculated for the upper euphotic zone during the same cruise (Landry et al., 2025), as well as those from other oligotrophic regions (Landry and Kirchman, 2002; Paffenhöfer et al., 2007). The low Chl*a* biomass reflected scarce concentrations of inorganic nutrients in the upper 50 m (NO_3_^2-^: <0.01-0.03 μM, PO_4_^3-^: ∼0.03-0.12 μM; Kranz et al., this issue). In the eastern Indian Ocean, a moderate fraction of C fixation can, in fact, originate from diazotrophy (N_2_ fixation) (Kranz et al., this issue; Nieblas et al., 2014; Sato et al., 2022), and lateral advection of nutrients (Kehinde et al., 2023).

Prokaryotes comprised almost 50% of microbial community carbon (C), and *Pro* was responsible for the Chl*a* biomass change observed across cycles and stations (linear regression, *p=*0.0226, not shown). Heterotrophic and mixotrophic nanoplankton dominated in abundance and biomass for the eukaryotic component. Given the small size and surface-to-volume ratio of the dominant cells and the prevalence of heterotrophs *vs* autotrophs, it is reasonable to think that these groups were selected for in these conditions (Chisholm, 1992), in contrast to productive upwelling areas dominated by larger autotrophs (e.g. diatoms). Mixotrophs and heterotrophs were key players in the eukaryotic community (Selph et al., this issue a) as estimated by flow cytometry and LTG staining, given the scarcity of inorganic nutrients and the rich prey size spectrum available (from bacteria to nanoplankton). Nutrient shortage is known to favor mixotrophy in marine plankton (Christaki et al., 1999; Hartmann et al., 2012; Unrein et al., 2014), enabling them to acquire organic nutrients through direct predation, symbiosis, and kleptoplastidy under conditions where obligate autotrophs would be limited (Flynn et al., 2019). Molecular analyses and high-performance liquid chromatography (HPLC) also confirm a high abundance of potentially mixotrophic taxa (Selph et al., this issue b) The abundant and diversified prey size spectrum could explain the higher abundance of nanoheterotrophs over mixotrophs, with their mainly higher ingestion and growth rates compared to non-specialist phagotrophs (Edwards et al., 2023; Jeong et al., 2005). Although LTG staining has been validated as a useful tool in previous studies (Anderson et al., 2018), this approach needs to be taken with caution given the potential staining of diatom silica deposition vesicles (Wilken et al., 2019). Yet, diatoms represented a negligible fraction of the protistan community in this region (Selph et al., this issue b), reducing the likelihood of non-specific LTG labelling (Table 4).

**Table 4.** Summary of research questions, pros and cons and potential biases of the methods applied and possible improvement strategies.

| Question | Method | Advantages | Disadvantages / Biases | Improvement |
| --- | --- | --- | --- | --- |
| How many cells of each population are present? | Flow cytometry | - Quali-quantitative (abundance, size, trophy)<br>- Short runs of very small cells<br>- Prokaryotes and eukaryotes<br>- Can be coupled to fluorescent stains (Hoechst, SybrGreen, LTG etc.) | - Poor taxonomic resolution<br>- Possible overestimation of trophic mode contributions due to non-specific labelling of LTG<br>- Underestimates fragile/large cells | - Ensure low diatom abundance to reduce non-specific LTG labelling<br>- Normalize the LTG+ signal stain against the LTG background noise<br>- Cross-calibrate with microscopy<br>- Apply gentle fixatives |
| How many cells do zooplankton eat? | Prey removal | - Established method especially with cultured prey<br>- Separates microzooplankton and copepod grazing<br>- Especially suited for selectivity estimates | - Internal grazing in mixed communities<br>- Requires assumptive corrections<br>- Bottle effect may under/over-estimate the real grazing impact (unrealistic patchiness) | - keep control bottles without predators to account for different grazer types<br>- Inoculate a reasonable number of copepods to avoid stress/competition<br>- Inoculation of nutrients to minimize effects of excretion by consumers |
| What type of diet do copepods rely on? | Isotope-labelling | - Avoids cell counts<br>- Detectable transfer across trophic steps<br>- Indicates assimilated nutrients rather than simple ingestion | - Non-specific isotope labelling<br>- Combined mixotroph and heterotroph contribution (both can consume labelled prey)<br>- Costly and sensitive to contamination<br>- Long incubations can dilute isotopic signal<br>- POM includes detritus | - Optimize incubation duration to balance signal vs. non-specific labelling<br>- Combine isotope tracers with other techniques, i.e., FACS or NanoSIMS<br>- Pair isotope data with microscopy/flow cytometry observations |

### 4.2. Copepod feeding: limitation by food quantity, quality and trophic cascades

Copepods cleared on average 75 mL ind^-1^ d^−1^, and ingested between 3.4 and 138 ng C ind^-1^ d^-1^ (median: 32.9 ng C ind^-1^ d^-1^), in agreement with field studies on copepod feeding on a natural plankton assemblage in oligotrophic conditions (2-400 mL ind^-1^ d^-1^, 10-200 ng C ind^-1^ d^-1^) (Broglio et al., 2004; Calbet and Saiz, 2005). The entire bottle volumes (108%) were cleared by the copepods, indicating perhaps that prey availability was not sufficient to sustain copepod demand. Occasionally, gut Chl*a* was also determined for the copepods (Exps 2, 8, and 9) and accounted for an equivalent of 0.8±0.4 µg Chl*a* ind^−1^ d^−1^ (for full description of the methodological procedure, see Décima et al., this issue). Hence, although our food removal method could not quantify copepod Chl*a* grazing, gut content data suggested recent ingestion equivalent to 32 ng C ind^-1^ d^-1^, when applying a C:Chl=58 as done for the integrated mesozooplankton community (Décima et al., this issue). Interestingly, some pelagic tunicates (appendicularians) were present in the incubation water. Copepods may have gained indirect access to Chl*a* especially from picoplankton trapped in the appendicularians’ mucous houses, given the ability of certain copepods to directly feed on them (Ohtsuka et al., 1996). Community grazing was also assessed via Chl*a* gut content analyses of size-fractionated net collections during the cruise and showed positive ingestion of Chl*a* by mesozooplankton (>0.2 mm) and up to 8% removal of Chl*a* production, though the highest Chl*a* grazing impact was exerted by zooplankton size fractions <1 and <0.5 mm (Décima et al., this issue).

It is possible that copepods fed on different prey that were not explicitly accounted for, such as large dinoflagellates and ciliates, known to comprise on average 33% and 43% of the diet in copepods in low nutrient environments (Kleppel, 1993; Saiz and Calbet, 2011). The highest contributions to dietary C were from nanomixotrophs and nanoheterotrophs, which are normally dinoflagellates and ciliate groups, preferred prey for copepods given their nutritional value and detectability (Saiz and Calbet, 2011). Despite some variability between experiments (e.g., Cycle 2, Transit), the overall dietary composition remained approximately stable. Different diet in Cycle 2 might be due to changes experienced by the community after a recent storm occurring at the end of Cycle 1 (Landry et al., 2025), yet hard to fully corroborate causality due to the limited amount of data. Whereas higher C intake in Transit experiment could be associated to a lower number of copepods used in the incubation (10 vs 50-70), increasing the individual C intake. Regarding the copepod grazing impact, other experiments also report enhanced grazing and selection on heterotrophic protists, but only a minimal impact on total production (2%) (Broglio et al., 2004), consistent with our observed nanoheterotroph removal rates (7.6%). It may also be true that larger prey might have contributed to high but undetected C intake. In fact, the dominant copepods sorted for incubation were mostly carnivores/omnivores and detritivores (*Candacia* spp., *Corycaeus* spp., *Euchaeta* spp., *Labidocera* spp., *Oncaea* spp., *Sapphirina* spp.), which preferentially feed on larger prey such as microplanktonic dinoflagellates, ciliates, copepod nauplii and marine snow (Atienza et al., 2006; Landry, 1978; Yen, 1985). Larger motile cells generate a stronger hydrodynamic disturbance making them easier to detect by rheotactic predators (Almeda et al., 2018). In addition, several calanoid copepods are unable to retain particles smaller than 5 µm due to larger distance between setae located on their mouthparts (Boyd, 1976). Due to these reasons, it is likely that pico- and small nanoplankton might have been suboptimal for efficient capture and retention by these copepods (∼1.5 mm prosome length).

However, bottle incubations are known to provide underestimates of absolute quantities of C consumed (Zeldis and Décima, 2020) and their best application is to investigate selectivity (Table 4). The underestimate is caused at least in part by the inability to replicate the local concentrations of prey encountered by a copepod. The ingestion calculation includes a clearance rate term (*F*, ml ind^-1^ d^-1^) and a concentration (cells mL^-1^), which is typically taken as the average concentration of the incubation or Niskin bottle, but likely does not represent small scale patchiness or thin layers exploited by copepods in the field to obtain their daily ration (Mullin and Brooks, 1976). In addition, low prey concentrations can be easily exhausted over the course of an experiment (Zeldis and Décima, 2020). However, when investigating selectivity of many types of prey, experiments need to be run long enough to detect the decrease or depletion of different preferred prey types. Thus, quantifying absolute ingestion and selectivity within a bottle experiment is usually not possible given the trade-offs described above, in addition to the inherent limitation of a bottle experiments not replicating the local prey concentration patchiness.

Copepods exhibit reduced selectivity under food limitation to increase the chance of food intake (Vincent and Hartmann, 2001). This and the high clearance rate observed might explain the absence of a clear preference for any of the five specific prey groups analyzed in our study, presumably in favor of other prey as previously pointed out. Though not significant, a moderate overall selection for mixotrophs emerged in parallel with avoidance of auto- and heterotrophs despite their abundance. Laboratory studies report no strong copepod preference based on trophic guild or nutritional value of their prey (Isari et al., 2013; Traboni et al., 2020). On the other hand, ingestion of phagotrophs can provide nitrogen, phosphorus and essential fatty acids, increasing the food quality for copepods (Klein Breteler et al., 1999) and enhancing the reproductive yield (Kleppel, 1993). Surprisingly, in our incubations, copepods seem to prefer and impact smaller picoplankton over nanoplankton. Compensatory feeding mechanisms might have led copepods to feed even on smaller prey to buffer the limiting food quantity (Burian et al., 2018). In a subtropical ecosystem, copepods fed natural nanoplankton (>2 µm, 2-5 µm, >5 µm) showed similar patterns of selection towards the smallest prey size class, ascribed to the dominance of picoplankton over other prey types, inducing copepods to maximize the intake of the most available prey (Calbet et al., 2000). Nevertheless, considering the large body size and the hunting behavior of the copepods in our experiments, active selection of picoplankton seems unlikely and our observations more likely the result of trophic cascades. This may have been triggered by microzooplankton preying on nanoplankton and releasing picoplankton from grazing pressure, which then became more abundant and more detectable and ingested by copepods. Microzooplankton grazing on nanoplankton may indirectly support pico-phytoplankton persistence, consistent with a trophic cascade. Nevertheless, the relative abundances of pico- and nano-sized prey are also shaped by intrinsic ecosystem structure, so a direct causal relationship cannot be confirmed from our experiments. Furthermore, appendicularians can compete with copepods for food and thus might have contributed to internal feeding together with protistan and metazoan micrograzers (Alldredge, 1984; Gorsky and Fenaux, 1998). As copepods ingested microzooplankton, the grazing pressure on Chl*a* standing stock would be released. Notably, Chl*a* concentration increased in copepod bottles by 30-60% compared to controls rather than declining due to grazing, supporting in part this cascade effect. However, one needs also to consider that by the end of the incubation (T_36_), the Chl*a* content of *Pro* cells (from flow cytometric measurements) increased (∼35%) significantly in the presence of copepods compared to bottles without copepods (*p*=0.037) in several experiments, suggesting that phototrophs might have benefitted from nutrients excreted by copepods, masking any copepod grazing impact on Chl*a*. This pinpoints the need, when possible, to add sufficient amount of inorganic nutrients to discard any effect of zooplankton excretion on phototrophic growth in food removal experiments (Table 4).

### 4.3. Microzooplankton grazing impact exceeds that of copepods

Microzooplankton were the dominant consumers of the microbial community, removing on average 110% of Chl*a* standing stock d^-1^, 126% of bacterial production and 111% of eukaryotic production. Grazing on prokaryotes removed 12% more production as on eukaryotes, less than the 33% difference reported by Landry et al. (2025) from in situ incubations in the upper euphotic zone. Our estimates differ as i) *Syn* and nanoheterotrophs were included only in this study, ii) our sampling was depth-specific, whereas Landry et al. (2025) sampled across a vertical gradient encompassing a wider range of conditions and determined rates over many more experiments, and iii) these incubations were done on deckboard incubators as opposed to in situ. Overall, our results align with the 40-100% range of daily primary production removal by microzooplankton in tropical and subtropical regions, including the Indian Ocean (Landry, 2009; Landry et al., 2022; Schmoker et al., 2013).

As expected for an oligotrophic ecosystem, copepods were less important grazers, removing on average 7% of protistan pico-nanoplankton production within the 2-12 µm size range. Consequently, the top-down control of eukaryotic production alone depended very little on copepods and was mostly driven by microzooplankton. Despite differences in grazing effort, both zooplankton groups consumed a high proportion of nanoheterotrophs, but copepods relied on nanomixotrophs to a higher extent.

### 4.4. Mixotrophy and intraguild predation: hidden pathways in oligotrophic food webs

Mixotrophs were important contributors to eukaryotic biomass. FlowCam images from the cruise detected putatively mixotrophic dinoflagellates (*Prorocentrum* spp., *Tripos* spp.) and ciliates (*Tontonia* spp.) (J. Goes, pers. comm.), as did 18S DNA sequences (Selph et al., this issue, b). A recent study in the same Indian Ocean region found rhizarians, including mixotrophic representatives, to be dominant together with copepods (Davies et al., 2022). Despite the absence of taxonomic detail for the mixotrophic and heterotrophic community in these experiments, LysoTracker Green coupled to flow cytometry (Selph et al., this issue a), and confirmed by HPLC and 18S data (Selph et al., this issue b) and previous community analysis (Davies et al., 2022) validate observations that mixotrophy is a widespread strategy spanning different protistan groups in this region, consistent with their well-known eco-evolutionary advantages in low-nutrient waters (Stoecker et al., 2017). Molecular analyses of microbial plankton from the eastern Indian Ocean have previously pointed to the elevated proportions of mixotrophic taxa, relative to declining contributions of true autotrophs and heterotrophs, in warm tropical waters that come from our study region (Raes et al., 2022). This also supports the hypothesis based on mesozooplankton grazing rates that the lower food web of these tropical waters may allow a more efficient trophic coupling to higher levels due to the high levels of mixotrophy (Landry et al., 2020a). While nanoheterotrophs contributed equally to what microzooplankton and copepod consumed in these experiments, nanomixotrophs were especially important for copepods. Phagotrophic protists provide important essential elements and can enhance nutrient transfer across trophic levels when ingested. Ecologically, this suggests that oligotrophic regions considered to be unproductive and not suitable for sustaining fisheries are actually a good nursery ground for fish larvae thanks to the compensation of nutrient depletion through mixotrophy and channeling, even if less efficient than the direct nutrient transfer scenario of rich ecosystems, and still allowing important species to thrive i.e., Southern Bluefin Tuna.

Copepod C ingestion rates estimated through the disappearance method were approximately an order of magnitude lower compared to those attained using stable isotope labelling. We can speculate on a few reasons for this discrepancy. From an ecological standpoint, we know that flow cytometry does not quantify the larger prey and consequently the disappearance estimates preclude the inclusion of their contribution to copepod diet. However, the contribution of microplankton to the overall plankton assemblage was very low (Yingling et al., this issue), such that it’s not likely that high enough abundances were present in our bottles to account for this difference. From a methodological standpoint, we used the δ^13^C values of POM as representative of what copepods were consuming. However, since POM includes all prey items and detritus (Stukel et al., 2014), it’s unlikely that copepods feed on all these components equally since copepods are highly selective (Broglio et al., 2004; Isari and Saiz, 2011). If the prey items consumed by copepods were small primary producers (with higher δ ^13^C values than POM because they are actively incorporating this tracer), then we have overestimated C-incorporation by using the values of POM. If the prey items consumed by copepods had lower δ^13^C values than POM because they were many trophic steps removed from primary producers, then our calculation underestimates C-consumption by copepods. The mean δ^13^C-based estimated C daily rations were 1.7% d^-1^, in the range of the estimates calculated for prey <15 µm in calanoid and cyclopoid copepods (Isari and Saiz, 2011; Schnetzer and Caron, 2005), but not sufficient to sustain basic respiratory and production rates at these temperatures (Décima et al., this issue; Landry et al., 2020b). However, these estimates are expected given the method, because bottles do not contain the prey densities to support the ingestion needed for respiration and production over 12 hours, as discussed above, but are in line with previous similar experiments (Isari and Saiz, 2011; Schnetzer and Caron, 2005).

When investigating patterns in the incorporation of N using the ^15^N-labelled prey, we cannot compare equally across cycles given the methodological differences associated with using *Syn* and *Pro*. However, we can compare patterns in ^15^N incorporation from Cycle 1 and 2 as representative of phagotrophic pathways connecting copepods to *Pro*, and Cycles 3, 4 and the transect, as representative of phagotrophic pathways connecting copepods to larger-sized prey similar to *Syn* (because *Syn* was not present in high abundances in these waters dominated by *Pro*). N daily rations were similar to C daily rations when we used *Syn* (both C and N approximately 1%), but were much lower when using *Pro* (1.8% C compared to 0.3% N). The higher N daily rations observed in *Syn* experiments point to a more efficient trophic pathway via the consumption of mixo- and heterotrophs (nano- and micro-sized), as confirmed by food removal incubations, which had more likely fed directly on ^15^N-labelled *Syn* due to its larger size (>1 µm). This highlights their presumptive role in nutrient channeling and trophic upgrading as previously reported for protozooplankton and more recently for mixoplankton (Klein Breteler et al., 1999; Ptacnik et al., 2004; Traboni et al., 2021). Selph and colleagues highlighted that mixotrophs were correlated with *Syn* during the cruise (Selph et al., this issue a), contributing to a high N transfer from ^15^N-*Syn* to copepods. We know that copepods did not feed directly on ^15^N-labelled *Syn* cells because the ^15^N-labelled prey had already been consumed by the time copepods were inoculated into the experimental bottles at T_24_. Regenerated nitrogen in the form of ammonia (NH_4_^+^) can be released by consumers of ^1^□N-labelled, and phytoplankton are capable of assimilating this pool on short timescales (Table 4). Yet, although plausible, this sequence is less likely to dominate over short incubations than direct ingestion of microbial consumers fed ^15^N prey, minimizing the impact of this indirect route to explain the observed copepod 1□N. We cannot rule out that copepods may have acquired 1□N by ingesting clumps of labelled cells that did not resuspend as single cells and/or that had been concentrated by appendicularians. However, appendicularians need to be collected very gently in order for them to exhibit natural behavior in containers (Masunaga et al., 2020; Scheinberg et al., 2005), which we did not do since they would have been transferred to our bottles directly from the Niskin. And while their abundances were high in our study, the highest concentrations were 200 individuals m^-3^ (Davies et al., this issue), which means that on average there was zero or 1 appendicularian per bottle (0.54 individuals in 2.7L), unlikely to significantly affect these results. Carnivory of copepods on younger stages, such as copepod nauplii and copepodites, which can feed on much smaller prey items than adult copepods, is potentially more likely given their higher concentrations (Davies et al., this issue; Swalethorp et al., this issue), and the inclusion of omnivorous/carnivorous taxa in our incubations. This could also explain the discrepancy in estimated C and N daily rations based on POM (which would include nauplii caught on the GF/F) and based on flow cytometry, which only includes direct consumption of unicellular prey.

In oligotrophic regions dominated by small-size cells, the ability of appendicularians to concentrate bacteria, picoplankton, and detritus into mucilaginous houses that they regularly abandon provides a major mechanism for transferring seemingly inaccessible nutrition to copepods. Copepods of the genus *Oncaea* are known to feed on marine snow and appendicularian houses (Ohtsuka et al., 1996) and were rather common in the Argo Basin community (Davies et al., this issue; Swalethorp et al., this issue). Appendicularians were also highly selected by *T. maccoyii* larvae during this cruise and accounted for up to 75% of their prey diet (Swalethorp et al., this issue), highlighting the role of gelatinous zooplankton in ichthyoplankton nutrition (Laiz-Carrion et al., 2015; Mianzan et al., 2001) and in nutrient transfer throughout the food web.

## 5. Summary and conclusions

Copepods grazed a small fraction of daily production, while microzooplankton are the main players in top-down control of protistan biomass in the Argo Basin. Mixotrophs and heterotrophs emerged as dominant functional groups in these oligotrophic food webs, supporting copepod and microzooplankton diets that channeled nutrients to higher trophic levels. Isotope tracing using labeled ^15^N-prey confirmed the importance of copepod consumption of phagotrophs and/or other metazoans, which was much more efficient at transferring energy from cells 1-2 µm compared to the <1µm-sized dominant *Prochlorococcus*. With ocean warming, smaller cell sizes (Morán et al., 2010) and increased phagotrophy (Wilken et al., 2013) are expected, making mixotrophy a competitive strategy for protist survival and increasing efficiency of energy transfer across trophic levels (Ward and Follows, 2016). This may help counteract the declining food quality predicted under nutrient limitation as a consequence of water-column stratification (Schenone et al., 2024). In regions dominated by small cells of poor stoichiometric and biochemical quality, copepod nutrition may strongly rely on larger, more nutritional prey, and benefit from the nutrient channeling and buffering mediated by microzooplankton and mixotrophs. The limitations of this study highlight the need for further investigations of prey diversity and selection towards or against specific taxonomic groups that encompass a larger size spectrum than the one examined herein.

## Declaration of competing interest

The authors declare no competing financial nor personal interests.

## Author statement

M.D. and C.T. conceived the study. C.T., G.F.C., K.E.S. and M.D. conducted sampling, experiments and sample processing onboard. K.E.S. processed flow cytometry data, G.F.C. processed samples for elemental content estimates. M.D. supported with analysis and interpretation of isotope data. K.E.S and M.R.L. provided cell biomass data, and support for interpretation of the results. C.T. analyzed results, conducted statistical analyses and drafted the manuscript. All authors contributed to provide conceptual feedbacks, comments, edits and thorough review of the manuscript.

## Acknowledgements

We are grateful to captain and crew of R/V *Roger Revelle* and to all the RR2201 science party. We thank N. Yingling and M. Stukel for providing stable isotope stocks, R. Swalethorp and C. Davies for assistance during zooplankton sampling, and J. Goes for providing FlowCam images of the protistan community. This study was supported by U. S. National Science Foundation grants OCE-1851558 (M.R.L.) and -1851381 (K.E.S.) and is a contribution to the Second International Indian Ocean Expedition (IIOE-2 endorsed project EP046). Seawater and plankton samples were collected under Australian Government permit AU-COM2021-520 and Australian Marine Parks permit PA2021-00062-2 issued by the Director of National Parks, Australia. Views expressed in this publication do not necessarily represent those of the Director of National Parks or the Australian Government. We thank two anonymous reviewers for their comments and suggestions that improved the quality and clarity of the manuscript.

## Declaration of generative AI and AI-assisted technologies in the manuscript preparation process

During the preparation of this work the author(s) used ChatGPT in order to improve the English and style of the text. After using this tool/service, the author(s) reviewed and edited the content as needed and take(s) full responsibility for the content of the published article.

## Notes

### Competing Interest Statement

The authors have declared no competing interest.

### Summary of Updates

The methodological limitations have been discussed in the new Table 4, the robustness of our approach and interpretation have been reinforced by literature evidence and independent data stemming from concurrent studies in the Argo Basin. The flow of the narrative has been improved.

